# STARMAP: A 3D-informed framework for mapping functional regions in proteins to regulatory and cellular phenotypes

**DOI:** 10.64898/2026.05.05.723010

**Authors:** Kriti Shukla, Jonnathan Castro, Darby Cheng, Lucy Holley, Elizabeth Brunk

## Abstract

Artificial Intelligence (AI) has transformed biology by revealing patterns in large-scale datasets and predicting regulatory relationships. Yet even the most advanced models often fail to identify biologically meaningful mechanisms from statistical associations. This limitation arises not from algorithmic capacity but from the lack of mechanistically grounded input features. Our structure-informed framework Structure-based Topological Analysis of Regulatory and Molecular Activity Patterns (STARMAP) embeds protein three-dimensional structure and population-scale functional genomics data into a unified representation for mechanistic inference. By mapping over 1.5 million naturally occurring variants across ∼1,700 cancer cell lines onto protein structures, STARMAP was able to identify spatial clusters of variation associated with shifts in transcriptional regulatory networks and drug response phenotypes. This approach transforms natural genetic variation into a large-scale, structure-informed screen, enabling systematic discovery of regulatory relationships across the proteome and providing interpretable and testable models of cellular regulation.

---

One of the central challenges in modern biology is no longer the lack of data, but how to organize large, complex datasets into mechanistic and testable insights^1^. Artificial Intelligence (AI) has greatly expanded our ability to detect patterns in high-dimensional biological data, yet most models still struggle to translate statistical associations into interpretable biological mechanisms. In many domains where AI performs well, the relationships of interest are directly represented in the data, for example in user interaction networks where structure can be inferred from observed connections. In contrast, biological measurements capture downstream, context-dependent consequences of regulatory activity rather than the underlying molecular interactions themselves.

A key limitation is therefore not the lack of data, but the lack of mechanistically meaningful representations^2,3^. Many gene regulatory network prediction approaches treat genes as independent features^4,5^ without incorporating the physical and functional context in which biological regulation occurs. Without inputs that reflect how regulation is encoded, models remain statistical black boxes. Here, we address this challenge by introducing a fundamentally different approach: embedding protein structure into large-scale functional genomics data. Protein structure provides a natural foundation for such representations because it defines how local perturbations give rise to shared functional consequences^6,7^. Within a protein, mutations that localize to the same structural region can perturb a common functional domain or interaction interface, leading to consistent downstream effects on interacting proteins and regulatory pathways^8^. For example, distinct mutations within a protein-protein interaction interface can each alter binding to a transcription factor, producing similar changes in its target gene program^8,9^.

To extend this principle to population-scale datasets, we developed a structure-informed framework that re-centers protein architecture as an organizing principle for mechanism discovery at scale. We call this framework Structure-based Topological Analysis of Regulatory and Molecular Activity Patterns (STARMAP). By organizing variants according to spatial proximity within protein structures, STARMAP captures shared structural perturbations and links them to shifts in transcriptional regulatory networks (TRNs) and drug response phenotypes. Leveraging matched population-scale genetic variation^10^, transcriptome^10–12^ and drug profiling^13^ data, together with AlphaFold structures^14^, STARMAP maps over 1.5 million naturally occurring variants onto protein structures and applies spatial statistical enrichment and deep learning to resolve coordinated regulatory and pharmacologic effects. Across nearly 15,000 proteins, this framework identifies more than 8 million regions of functional influence that act as focal points of system-level behavior.

More broadly, STARMAP demonstrates that embedding biophysical context into large-scale datasets enables interpretable models of variant effects and drug sensitivity. This framework enables proteome-wide discovery of structure-function relationships that are inaccessible to existing approaches. To facilitate exploration and reuse, we provide an interactive resource that allows users to query structure-resolved associations between protein regions, regulatory programs, and drug responses across the proteome (available at: starmap.unc.edu).

## RESULTS

### STARMAP framework for structure-aware mapping of molecular perturbations

To systematically link genetic variation to cellular phenotypes, we assembled a matched population-scale multi-omics resource that spans ∼1,700 cancer cell lines^10–12^ and integrates large-scale genomic variation^10^, bulk transcriptomic profiles^10–12^, pharmacologic response data^13^, AlphaFold structures^14,15^ and single-cell perturbation data^16–18^ (***Fig. 1a, Extended Data 1a***). Aggregated across cell lines, mutation data define a large-scale “variome” comprising over one million naturally occurring variants and enabling a systematic analysis of genotype-phenotype relationships (***Fig. 1b, Extended Data 1b***). Mapping these variants onto three-dimensional protein structures reveals that mutations can cluster in spatial neighborhoods, even when they are distant in sequence or rare in frequency (***Fig. 1c, Extended Data 1c***), uncovering functional relationships that are not apparent in linear representations^7,19–21^.

**Figure 1.**
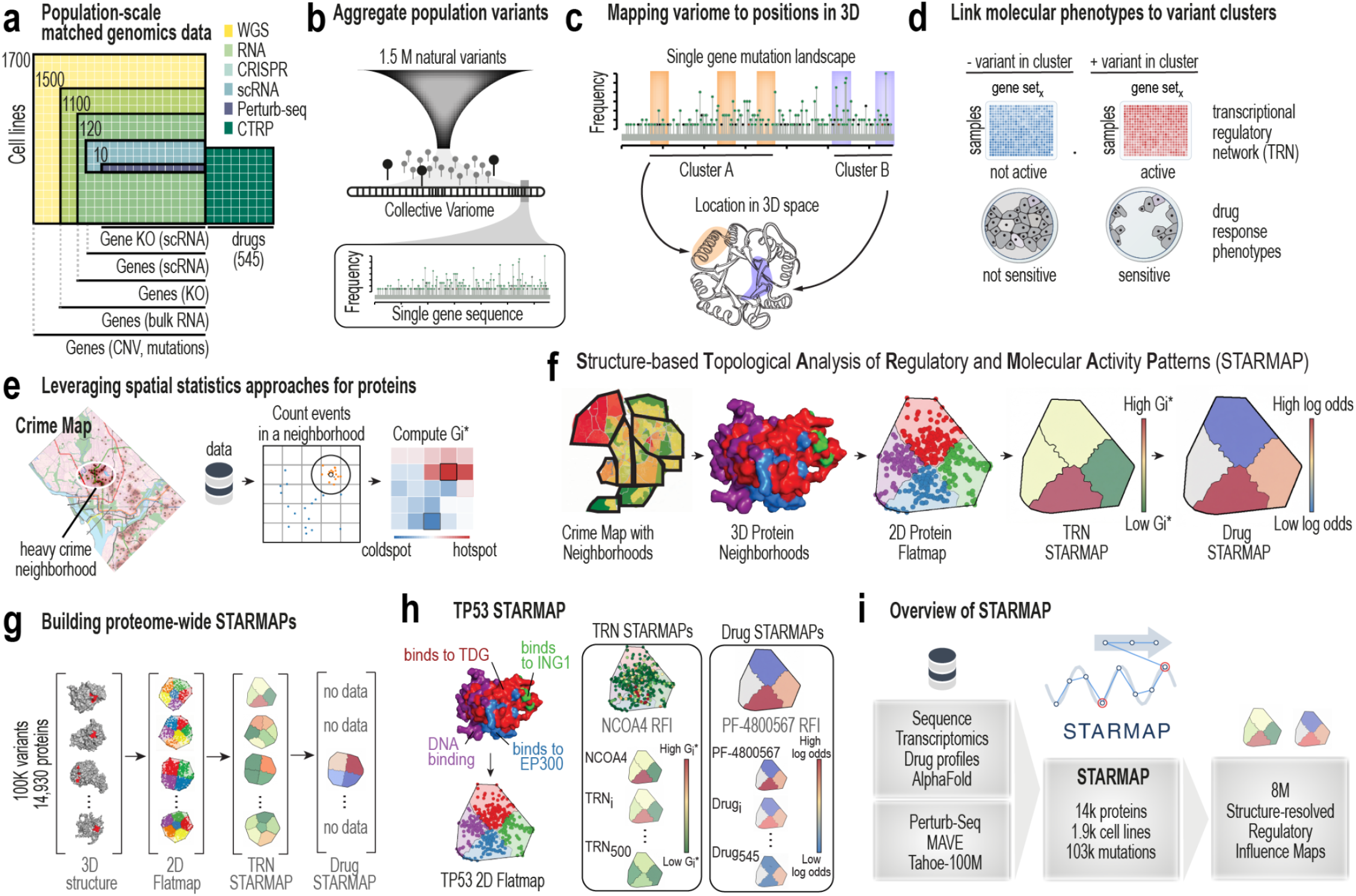
STARMAP framework for structure-aware mapping of molecular perturbations. **a**, Overview of matched population-scale multi-omics data across ∼1,700 cancer cell lines, integrating genomic variation (whole genome sequencing -WGS), transcriptomic profiles (RNA), perturbation screens (CRISPR), and drug response measurements. **b**, Representation of the population-scale variome, comprising naturally occurring variants across the proteome. **c**, Mapping of variants from linear sequence space onto three-dimensional protein structures, revealing spatial clustering independent of sequence position or mutation frequency. **d**, Conceptual framework for associating structurally localized variant clusters with shared phenotypic outcomes across cell lines, including TRN activity and drug response. **e**, Regions enriched for events in the protein structures are identified using a spatial statistics method employed for geographic hotspot (in this example, crime) detection. This method allows for calculation of the TRN association score (Gi*). **f**, STARMAP workflow: protein structures are partitioned into local neighborhoods and projected onto two-dimensional flatmaps for visualization; spatial statistics identify regions of functional influence associated with phenotypic signals. **g**, Proteome-wide generation of TRN-specific and drug-specific STARMAPs, linking protein regions to regulatory and pharmacologic phenotypes. **h**, Example STARMAP for TP53, highlighting structural regions associated with specific transcriptional programs and drug responses. **i**, Summary of STARMAP inputs and outputs, including integrated multi-omics data and millions of structure-resolved regulatory influence maps across the proteome.

STARMAP groups variants within each protein based on spatial proximity in three-dimensional structure and tests whether these localized clusters are associated with shared phenotypic outcomes across cell lines ***(Fig. 1d)***. Specifically, we evaluate whether variants within a structural region are associated with shifts in transcriptional regulatory network (TRN) activity or drug response. To quantify these associations, we adapt spatial statistics from geographic analysis to the protein structural context. In geographic applications, these methods are used to identify hotspots of activity, such as regions within a city where incidents of crime occur more frequently than expected ***(Fig. 1e)***. Analogously, we construct a “STARMAP” for each protein, in which the three-dimensional structure is represented as a two-dimensional flatmap for visualization, and regions are colored according to the strength of their association with transcriptional or pharmacologic changes ***(Extended Data 1d)***. In this representation, hotspots correspond to regions of transcriptional or drug influence, where variants are disproportionately associated with coordinated phenotypic shifts, while coldspots represent regions with minimal or no association. Proteins are partitioned into local structural neighborhoods, and a modified Getis-Ord Gi* statistic^22^ is used to detect regions enriched for TRN activity shifts. Regions with high Gi* scores are defined as <u>R</u>egions of <u>T</u>ranscriptional <u>I</u>nfluence (RTIs). In parallel, a modified log-odds statistic is used to identify regions associated with changes in drug sensitivity, termed <u>R</u>egions of <u>D</u>rug Influence (RDIs). Together, RTIs and RDIs define structural hotspots where localized perturbations are linked to system-level regulatory and pharmacologic behavior (***Fig. 1f, Extended Data 1e***).

Applying the STARMAP framework across proteins enables construction of structure-resolved maps linking protein regions to regulatory and pharmacologic phenotypes (***Fig. 1g***).For each protein, STARMAP generates hundreds of TRN-specific and drug-specific influence maps. For example, analysis of TP53 reveals distinct structural regions associated with specific regulatory programs and drug responses, many of which correspond to known functional interfaces and interactions (***Fig. 1h***). Across the proteome, STARMAP generates more than 8 million structure-resolved associations spanning over 14,000 proteins, 500 TRNs, and 545 drugs, providing a scalable framework for linking genetic variation to cellular phenotypes (***Fig. 1i, Extended Data 1f***). To enable exploration and reuse of these data, we provide an interactive resource that allows users to query structure-resolved associations across proteins, regulatory networks, and drug responses (***Extended Data 1g, 2, 3***; starmap.unc.edu).

### Structural regions define transcriptional regulatory programs

To determine whether specific regions within proteins are associated with gene regulatory networks, we constructed transcriptional regulatory network (TRN) STARMAPs for all proteins with genomic variation data in DepMap^10–12^ (***Fig. 2a-b***). Variants were mapped onto AlphaFold structures^14^ and evaluated for association with the activity of more than 500 TRNs, quantified using Gene Set Enrichment Analysis^23,24^ on transcription factor-target gene sets^25,26^.

**Figure 2.**
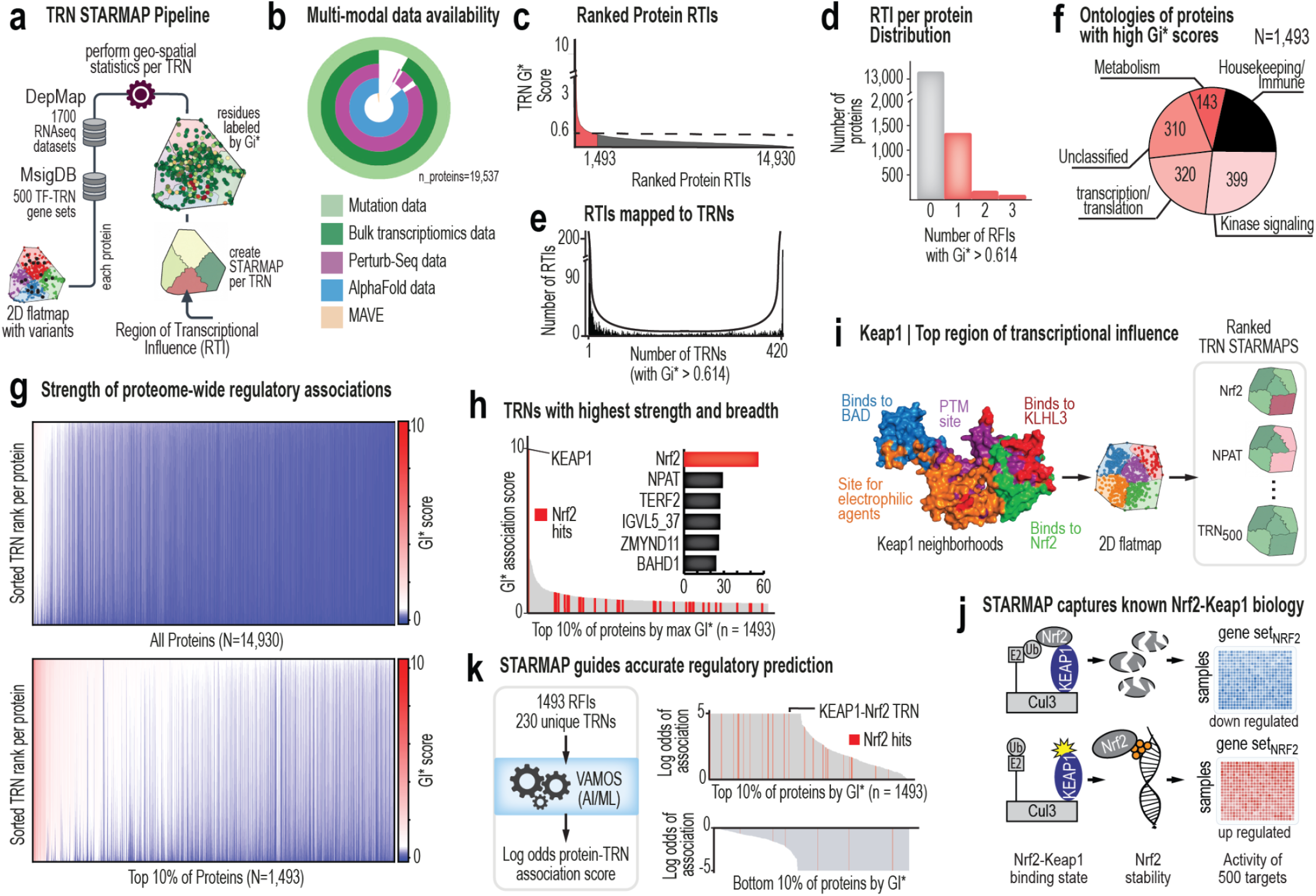
Structural regions define transcriptional regulatory programs. **a**, Computational framework for generating TRN STARMAPs, mapping variants onto protein structures and identifying regions associated with TRN activity. **b**, Coverage of multi-modal datasets used to construct STARMAPs, including genomic variation, transcriptomics, and perturbation-based measurements. **c**, Distribution of TRN association scores (Gi*) across proteins, highlighting a subset with strong regional associations. **d**, Number of RTIs per protein, showing that most proteins contain few or no RTIs. **e**, Distribution of TRN associations per protein, revealing a bimodal pattern of regulatory specificity versus pleiotropy. **f**, Functional annotation of proteins with strong TRN associations, showing enrichment for regulatory and signaling roles. **g**, Global visualization of RTIs across proteins, illustrating sparsity and heterogeneity of structure-regulatory associations. **h**, Top TRN associations across proteins, highlighting NRF2 as a dominant regulatory signal. **i**, STARMAP of KEAP1, identifying a hotspot of regulatory influence within the NRF2-binding interface. **j**, Schematic of KEAP1-NRF2 regulatory mechanism and its disruption by mutations. **k**, Predictive modeling of TRN activity using STARMAP-defined RTIs and unique TRNs, demonstrating high-confidence associations.

For each protein, spatial statistics were used to identify regions associated with significant shifts in TRN activity. Specifically, we assessed whether cell lines harboring variants within a shared structural region are enriched for higher or lower target gene expression relative to the population. Regions with high TRN association scores (Gi*) (termed Regions of Transcriptional Influence or RTIs) represent structural loci statistically linked to a specific transcriptional program. Despite the large search space, exceeding seven million possible associations, strong structure-regulatory relationships are rare (***Fig. 2c***). Most proteins contain no RTIs or a single RTI, with only a small subset exhibiting multiple regions of transcriptional influence (***Fig. 2d***). Proteins tend to fall into two distinct classes: those associated with a single TRN (specific) and those linked to many TRNs (pleiotropic), with relatively few intermediate cases (***Fig. 2e***). Importantly, TRN association scores show consistent scaling and are not driven by protein length or mutation frequency ***(Extended Data 4a–f)***, supporting the robustness of the approach.

Proteins with the strongest TRN associations are enriched for signaling, regulatory, and chromatin-associated functions ***(Fig. 2f)***. Notably, a substantial fraction of these proteins lack prior functional annotation, indicating that STARMAP captures both known and previously uncharacterized regulators of transcription. Global visualization of RTIs further highlights the sparsity and heterogeneity of these associations, with only a small subset of regions exhibiting strong regulatory influence ***(Fig. 2g)***.

Among the most prominent signals, NRF2-mediated transcription emerges as a dominant and recurrent regulatory program across proteins ***(Fig. 2h)***. The strongest association is observed for KEAP1, a well-characterized regulator of NRF2 stability^27–30^ (***Fig. 2i, Extended Data 4g***). STARMAP identifies a hotspot within the NRF2-binding interface of KEAP1^29^, consistent with its known role in targeting NRF2 for degradation^30,31^. Mutations in this region are associated with increased NRF2 activity^8,32,33^, reflecting disruption of KEAP1-mediated repression (***Fig. 2j***). To assess predictive power, we applied a machine learning framework (VAMOS^8,9^) to predict NRF2 target gene activity from natural variation in DepMap cell lines. Models trained on STARMAP-defined regions achieved high predictive performance, with the KEAP1-NRF2 interaction among the strongest and most confident associations (Log odds = 5.000, ***Fig. 2k***).

### Independent validation of structure-regulatory associations using single-cell perturbation data

To independently validate STARMAP-derived structure-regulatory associations, we leveraged the X-Atlas Orion single-cell Perturb-seq dataset^16^ that directly measures transcriptional responses to targeted genetic perturbations^17^ (***Fig. 3a, Extended Data 5a***). For each cell, TRN activity was quantified using gene set enrichment analysis across transcription factor target gene sets25,26 and compared to STARMAP-predicted associations. Specifically, we tested whether perturbation of genes linked to a given structural region produces concordant changes in the corresponding TRN activity, providing an orthogonal validation of structure-regulatory relationships..

**Figure 3.**
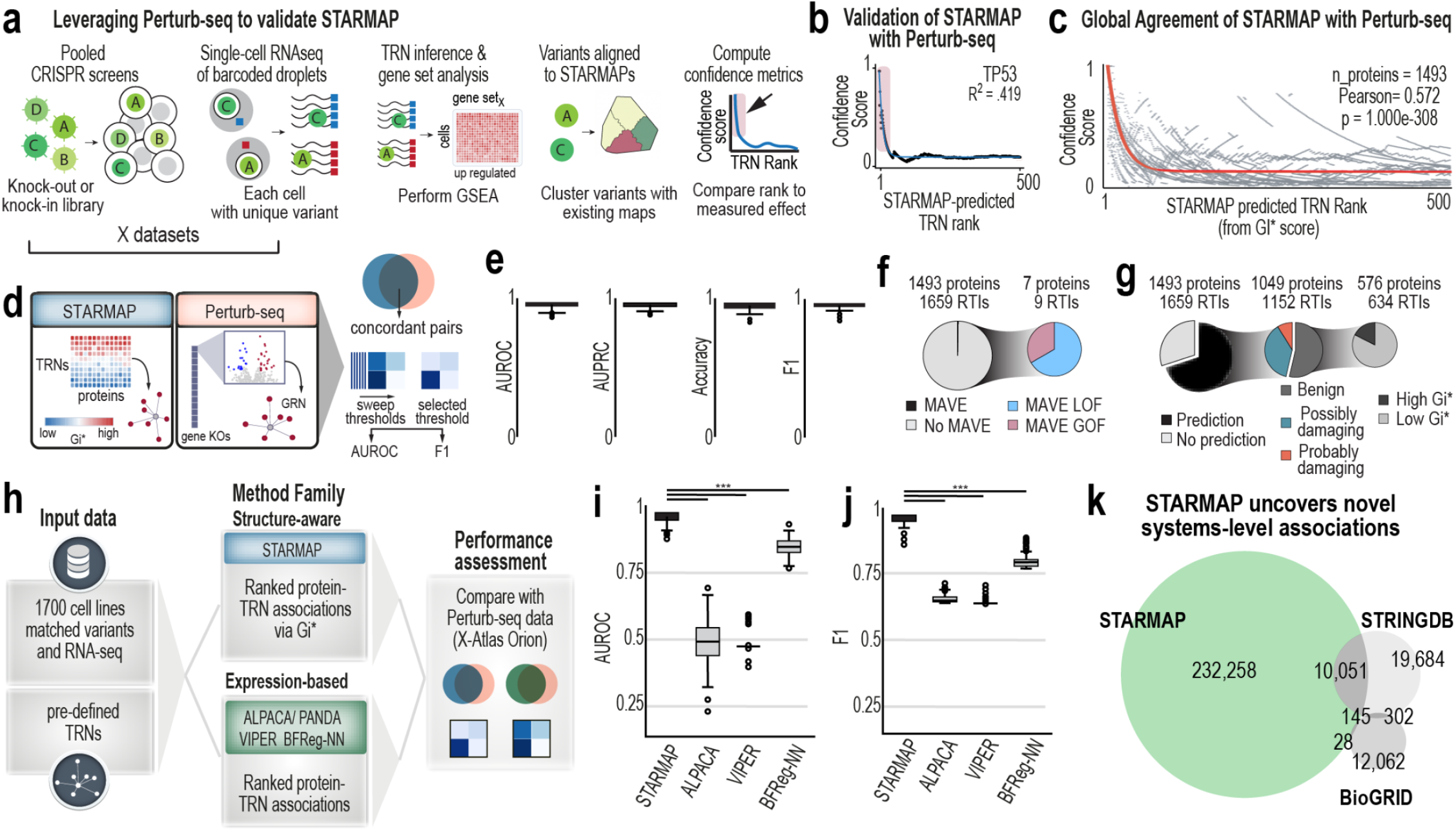
Independent validation of structure-regulatory associations using single-cell perturbation data. **a**, Overview of validation strategy using Perturb-seq. Pooled CRISPR perturbations were applied across cell populations, followed by single-cell RNA sequencing to measure transcriptional responses. TRN transcriptional activity in the form of a GSEA score is calculated per cell, enabling comparison between experimentally observed regulatory changes and STARMAP-predicted TRN association scores (GI*). **b**, Example validation for TP53. Confidence scores for Perturb-seq-derived TRN transcriptional activity are plotted against STARMAP-predicted TRN rank, derived from TRN association scores (GI*), showing strong enrichment of high-confidence signals among top-ranked TRNs. **c**, Global agreement between STARMAP predictions and Perturb-seq measurements across proteins and TRNs, demonstrating consistent alignment between predicted and observed regulatory activity (Pearson r ≈ 0.572). **d**, Schematic of validation framework. STARMAP-derived TRN rankings are compared to Perturb-seq-derived transcriptional responses by evaluating concordance between predicted regulatory associations and experimentally observed gene expression changes. **e**, Performance metrics quantifying agreement between STARMAP predictions and Perturb-seq data, including AUROC, AUPRC, F1 score, and accuracy. **f**, Comparison of STARMAP-identified regions with MAVE. **g**, Relationship between STARMAP-identified variants and PolyPhen predictions, showing that structure-based grouping captures both predicted damaging and nominally benign variants. **h-j**, Comparison of STARMAP with existing gene regulatory network inference methods. **k**, Concordance of STARMAP-derived associations with STRING database text-mining annotations and BioGRID known protein-protein interactions.

We first assessed the agreement between STARMAP-derived TRN rankings and Perturb-seq-derived regulatory responses. For individual proteins such as TP53, highly ranked TRNs identified by STARMAP show strong concordance with experimentally observed transcriptional responses, with top-ranked TRNs exhibiting consistently high confidence scores (***Fig. 3b, Extended Data 5b***). This relationship extends globally across proteins and regulatory networks, with STARMAP predictions showing strong agreement with Perturb-seq measurements (Pearson r ≈ 0.52), indicating that structure-defined regions capture biologically meaningful regulatory signals (***Fig. 3c, Extended Data 5c***). To quantify this agreement, we compared STARMAP-derived TRN rankings with Perturb-seq-derived transcriptional responses using a classification framework ***(Fig. 3d)***. For each protein, TRNs were ranked based on STARMAP association scores and evaluated against experimentally derived responses using rank-based confidence metrics and standard classification measures. This analysis demonstrates strong concordance between STARMAP predictions and experimentally observed perturbations at single-cell resolution ***(Fig. 3e)***.

We next assessed whether STARMAP-identified regions correspond to functional effects at the protein level. Comparison with multiplexed assays of variant effects^34–36^ shows that variants within RTIs are enriched for functional impact, with the majority exhibiting loss-or gain-of-function behavior ***(Fig. 3f)***. Similarly, integration with PolyPhen^6^ predictions indicates that STARMAP captures both predicted damaging and nominally benign variants ***(Fig. 3g)***, suggesting that structure-based grouping provides complementary information beyond sequence-based annotation.

Finally, we compared STARMAP to existing gene regulatory network inference methods, including expression-based^37^ and network-based methods^38–40^. STARMAP consistently outperforms these methods in identifying Perturb-seq-derived protein–TRN associations ***(Fig. 3h, Extended Data 5d–g)***, demonstrating improved ability to recover experimentally observed regulatory relationships. Comparison with protein-protein interaction databases, including STRING^41–52^ and BioGRID^53^, further shows that STARMAP captures both known interactions and a large set of previously uncharacterized regulatory associations (***Fig. 3k***).

### Structural mapping of drug responses reveals spatial determinants of pharmacologic sensitivity

To determine whether localized structural perturbations are associated with drug response, we developed a structure-informed machine learning framework that links variant position to pharmacologic sensitivity ***(Fig. 4a)***. For each protein, variants were mapped onto three-dimensional structures^14^, and spatial features, including cluster identity and distance to annotated functional sites, were used to train a multilayer perceptron model^54^ to predict drug sensitivity across cell lines using binarized response profiles from the Cancer Therapeutics Response Portal (CTRP)^13^.

**Figure 4.**
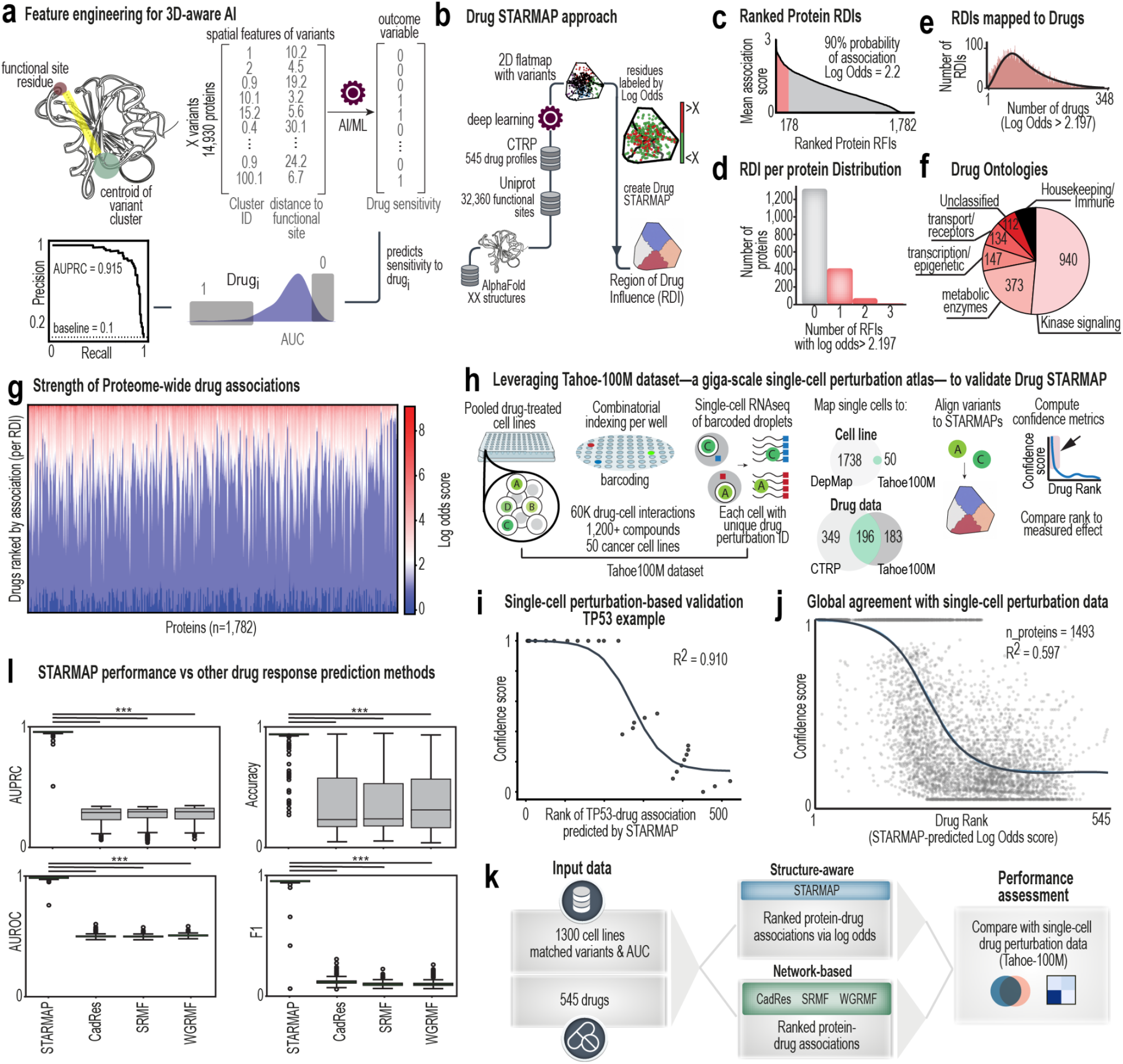
Structural mapping of drug response reveals spatial determinants of pharmacologic sensitivity. **a**, Structure-aware machine learning framework for predicting drug response. Variants were mapped onto protein structures, and spatial features, including cluster identity and distance to functional sites, were used to model drug sensitivity across cell lines. **b**, Variant-drug associations were mapped onto protein structures and aggregated within spatial neighborhoods to identify RDIs. **c**, Distribution of protein–drug sensitivity association scores (log-odds) with a subset of proteins with high-confidence associations highlighted (log odds > 2.2). **d**, Number of RDIs per protein. **e**, Distribution of drug associations per protein. **f**, Functional annotation of proteins with RDIs that have high drug sensitivity association scores, showing enrichment for kinase signaling, metabolic enzymes, and transcriptional regulators. **g**, Global visualization of protein–drug associations across the proteome, illustrating heterogeneity in drug sensitivity patterns. **h**, Overview of validation using Tahoe-100M single-cell drug perturbation data. **i**, Example validation for TP53, showing agreement between STARMAP-predicted drug associations and single-cell transcriptional responses. **j**, Global agreement between STARMAP predictions and single-cell perturbation data across proteins and drugs. **k–l**, Comparison of STARMAP performance with existing drug response prediction methods.

Building on these predictions, we constructed Drug STARMAPs by mapping model-derived association scores onto protein structures ***(Fig. 4b)***. For each protein-drug pair, variants were aggregated within structural neighborhoods and evaluated using log-odds statistics to quantify associations with drug sensitivity. RDIs (or Regions of Drug Influence) represent localized structural regions whose perturbation is strongly associated with drug response ***(Extended Data 6a)***.

Across the proteome, strong region-drug associations are sparse. Only a subset of proteins exhibit high-confidence RDIs (log odds > 2.2), corresponding to approximately 90% probability of association with a specific drug ***(Fig. 4c)***. Consistent with this, most proteins contain zero or one RDI, with only a small number exhibiting multiple RDIs ***(Fig. 4d)***. Drug associations are similarly uneven, with a small subset of compounds accounting for the majority of RDIs ***(Fig. 4e)***. Proteins with strong RDIs are enriched for signaling pathways, particularly kinase-associated proteins, as well as metabolic enzymes and transcriptional regulators ***(Fig. 4f)***. Global visualization further highlights the heterogeneity of protein-drug relationships, with distinct patterns of sensitivity across proteins and drugs ***(Fig. 4g, Extended Data 6b)***.

To validate these associations, we leveraged the Tahoe-100M single-cell perturbation dataset^18^ (***Extended Data 6c***), which measures transcriptional responses to drug treatment at single-cell resolution ***(Fig. 4h, Extended Data 6c)***. By aligning shared cell lines and drugs across datasets ***(Extended Data 6d)***, we compared STARMAP-predicted drug associations with experimentally observed transcriptional responses. For individual proteins, such as TP53, predicted associations show strong agreement with single-cell perturbation data, with high-confidence signals concentrated among top-ranked drug associations ***(Fig. 4i, Extended Data 6e)***. This relationship extends globally, with strong concordance observed across proteins and drugs ***(Fig. 4j, Extended Data 6f)***. Finally, we compared STARMAP to existing drug response prediction methods, including matrix factorization^55^ and similarity-based approaches^56,57^. Across all evaluation metrics, STARMAP demonstrates improved predictive accuracy and robustness relative to these methods (***Fig. 4k–l, Extended Data 6g-h***).

### Regulatory and pharmacologic perturbations occupy a structured network space

To understand how perturbations within specific protein regions propagate across cellular systems, we constructed protein-region-centric networks linking RTIs to TRNs ***(Fig. 5a, Extended Data 7a)***. In this representation, protein regions are connected based on shared TRN associations, revealing higher-order structure in regulatory space. Distinct protein regions converge onto common regulatory programs, forming interconnected modules of transcriptional control.

**Figure 5.**
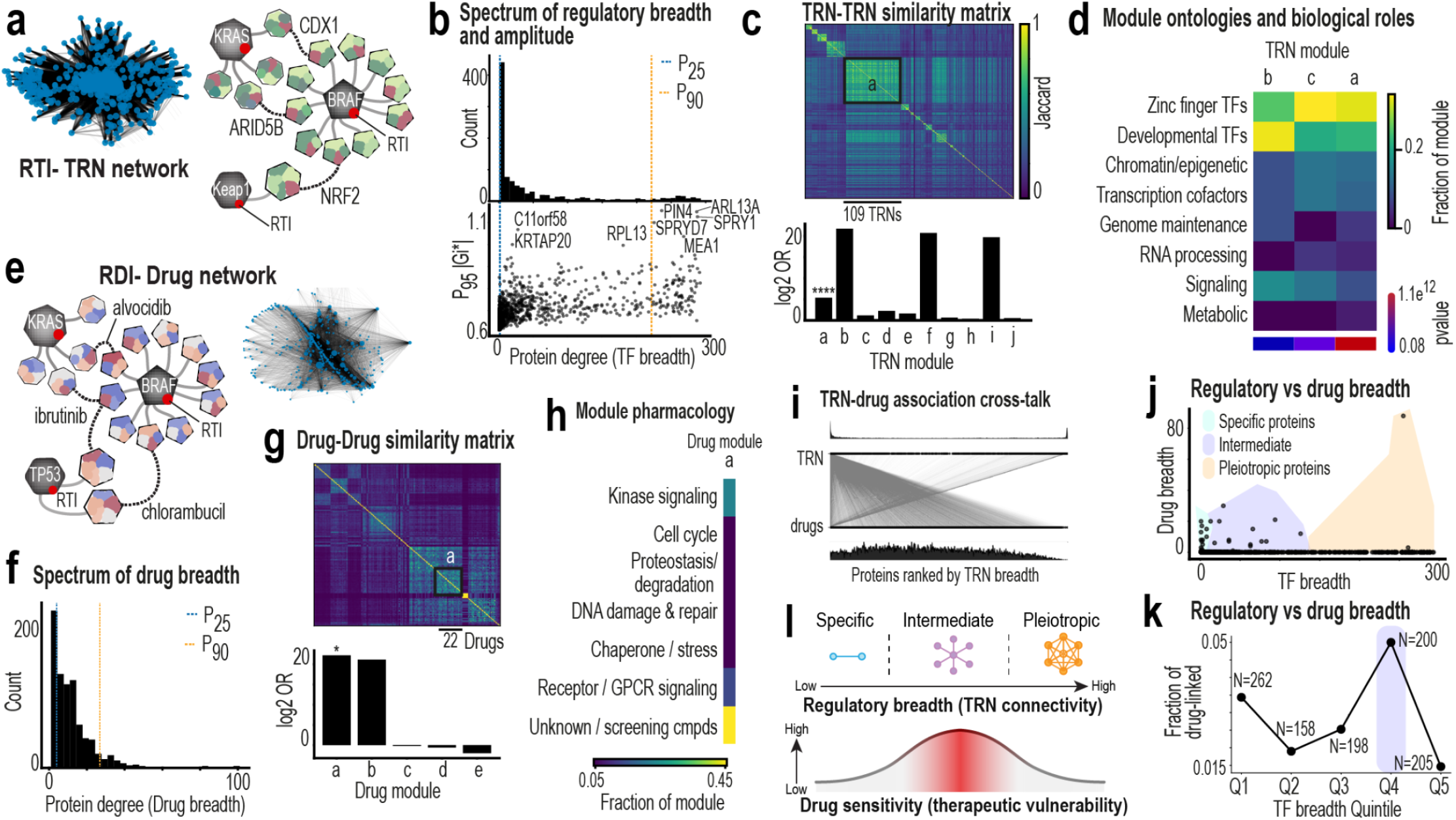
Regulatory and pharmacologic perturbations occupy a structured network space. **a**, Protein-centric network linking RTIs through shared TRNs. Example connections illustrate convergence of distinct protein regions onto common regulatory programs. **b**, Distribution of regulatory breadth (number of TRNs per protein) and regulatory amplitude (95th percentile TRN association score - GI*), showing distinct regimes of specific and pleiotropic regulatory influence. **c**, TRN-TRN similarity matrix based on shared protein associations, revealing modular organization of regulatory space. **d**, Functional annotation of TRN modules, highlighting enrichment for transcriptional, chromatin-associated, and regulatory processes. **e**, Protein-centric network linking RDIs based on shared drug associations. **f**, Distribution of drug breadth across proteins. **g**, Drug-drug similarity matrix based on shared protein associations, revealing modular pharmacologic structure. **h**, Functional annotation of drug modules, indicating enrichment for kinase signaling, cell-cycle regulation, stress response, and related pathways. **i**, Comparison of regulatory and pharmacologic associations across proteins, illustrating differences in connectivity between TRN and drug networks. **j**, Relationship between regulatory breadth and drug breadth, showing that drug associations are enriched in proteins with intermediate regulatory connectivity. **k**, Quantification of drug linkage across regulatory breadth quantiles, demonstrating peak drug sensitivity in intermediate regimes. **l**, Conceptual model illustrating the relationship between regulatory breadth and pharmacologic sensitivity, highlighting an intermediate regime of network engagement associated with therapeutic vulnerability.

Proteins exhibit a broad spectrum of regulatory breadth, defined by the number of TRNs to which each protein is linked, as well as regulatory amplitude, reflecting the strength of an association (***Fig. 5b, Extended Data 7b-g***). Two dominant regimes emerge: proteins with highly specific regulatory influence and those with broad, pleiotropic connectivity across many TRNs with relatively few proteins occupying intermediate states. Clustering TRNs based on shared protein associations reveals modular organization within regulatory space (***Fig. 5c***), with a subset of TRNs forming highly interconnected modules enriched for pleiotropic proteins. Functional annotation of these modules indicates enrichment for core regulatory processes, including transcriptional control and chromatin-associated functions (***Fig. 5d***).

We constructed analogous networks for pharmacologic response by linking RDIs across proteins ***(Fig. 5e)***. In contrast to regulatory networks, drug associations are more constrained, with most proteins exhibiting limited drug breadth ***(Fig. 5f)***. Drug-drug similarity analysis reveals modular organization of pharmacologic space, with modules corresponding to shared mechanisms of action, including kinase signaling, cell-cycle regulation, and stress response pathways ***(Fig. 5g-h)***. These patterns reflect both intrinsic protein biology and drug-specific effects rather than network structure alone ***(Extended Data 7f)***.

Comparison of regulatory and pharmacologic networks reveals a key distinction: proteins with broad regulatory influence are not necessarily associated with broad drug sensitivity ***(Fig. 5i)***. Instead, drug associations are enriched among proteins with intermediate regulatory breadth, while highly pleiotropic proteins show relatively limited pharmacologic linkage. Quantitative analysis confirms that drug sensitivity peaks within intermediate quantiles of regulatory connectivity ***(Fig. 5j-k)***. While many perturbations induce widespread regulatory effects, only a subset, those within an intermediate regime of network engagement, are strongly associated with drug sensitivity. This defines a constrained regime of network perturbation that is linked to therapeutic vulnerability and provides a principled framework for identifying actionable molecular states ***(Fig. 5l)***.

### Structural mapping of KRAS variants reveals mechanistic and clinical insights

To assess whether STARMAP can uncover mechanistic relationships, we focused on KRAS, a well-characterized oncogene^58–60^. Integration of population-scale perturbation datasets shows that KRAS is uniquely suited for multi-modal analysis, with matched variant^10,11^, transcriptional^10,12^, Perturb-seq^16^, and functional assay^36^ data available (***Fig. 6a***). STARMAP analysis identifies distinct regions of transcriptional and drug influence, with strong enrichment of specific TRNs and drug associations within a defined structural cluster ***(Fig. 6b–c)***. Among these, TEAD2-associated transcriptional activity emerges as a dominant regulatory signal linked to variants in this region. Mapping these associations onto the KRAS structure shows that both transcriptional and drug-associated hotspots localize to a functionally annotated interface, suggesting that perturbations in this region drive coordinated downstream effects (***Fig. 6d***). To interpret these associations, we integrated multi-omic data with prior biological knowledge to propose a mechanistic model linking KRAS variants to the TEAD2 transcriptional program^61,62^ ***(Fig. 6e)***. Variants within this cluster are associated with increased MAPK signaling activity^63–67^ and decreased Hippo pathway activity^68^, consistent with activation of YAP/TAZ-mediated transcription^69–74^. Accordingly, TEAD2-associated TRN activity is elevated in KRAS variant cell lines^75^. Drugs identified by STARMAP as associated with this region show effects consistent with this regulatory axis^76,77^. Treatment with dinaciclib or homoharringtonine suppresses TEAD2-associated activity specifically in KRAS variant contexts (***Fig. 6e***), suggesting that these compounds modulate pathways linked to the structural perturbation.

**Figure 6.**
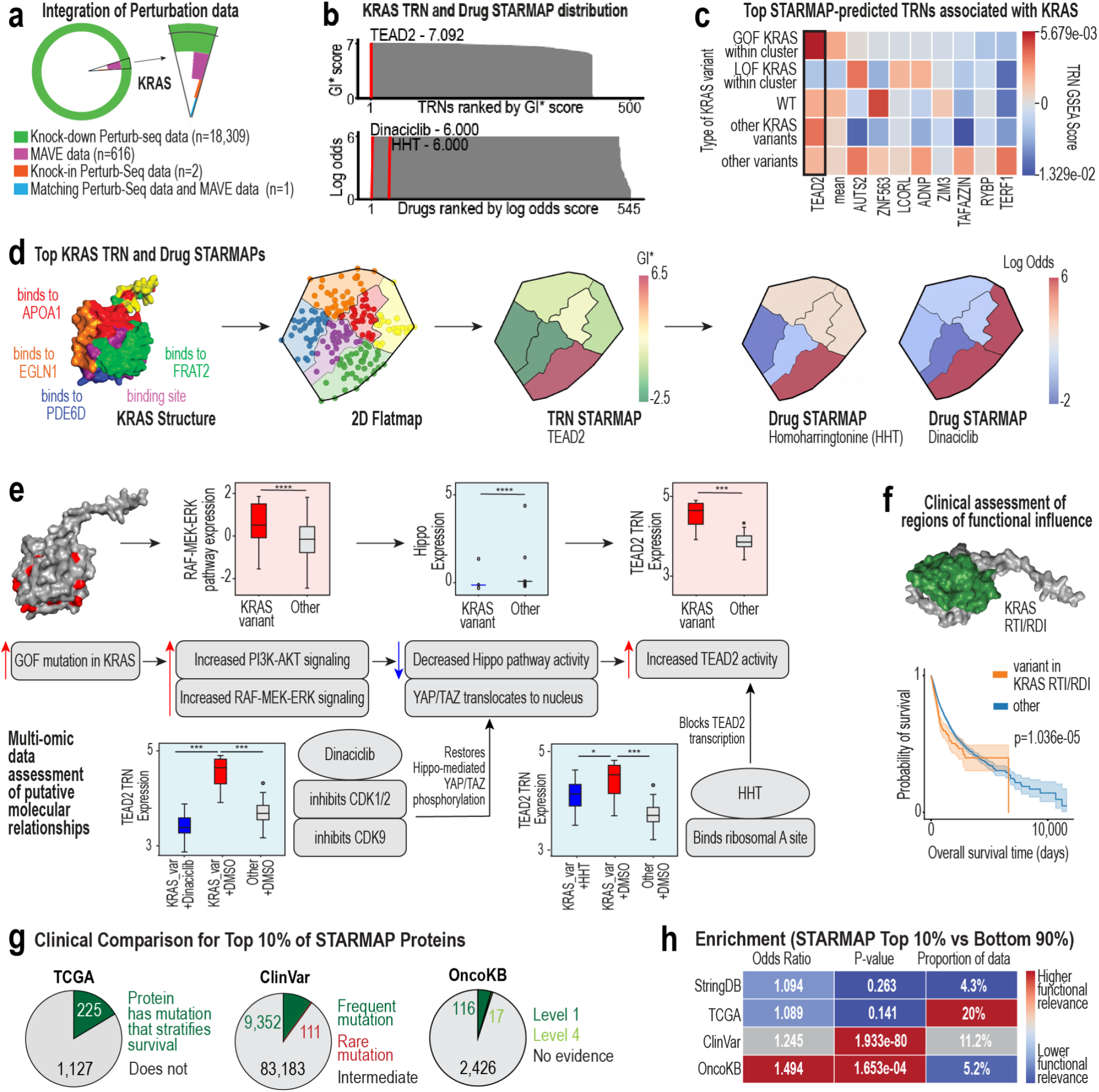
Structural mapping of KRAS variants reveals mechanistic and clinical insights. **a**, Integration of multi-modal perturbation datasets for KRAS, including variant, Perturb-seq, and functional assay data. **b**, Distribution of TRN association scores (Gi*) and drug sensitivity association scores (log-odds) across KRAS structural regions. **c**, Heatmap of TRN activity across KRAS variant classes, highlighting enrichment of TEAD2-associated transcriptional programs in specific variant clusters. **d**, Structural mapping of KRAS showing annotated functional regions, corresponding flatmap representation, and STARMAPs for TRN and drug associations, highlighting overlapping regions of influence. **e**, Proposed mechanistic model linking KRAS structural perturbations to downstream signaling and transcriptional programs. Transcriptomic validation of the proposed mechanism, showing increased MAPK signaling, decreased Hippo pathway activity, and elevated TEAD2-associated transcriptional activity in KRAS variant cell lines. Drug perturbation experiments demonstrating suppression of TEAD2-associated transcriptional activity in KRAS variant contexts following treatment with dinaciclib and homoharringtonine. **f**, Kaplan–Meier analysis showing reduced overall survival for patients with KRAS variants localized to the RTI or the RDI compared to other KRAS variants. **g**, Proteome-wide assessment of the top 10% of STARMAP proteins (TRN association score - Gi*>0.612, n=1,493) against TCGA, ClinVar, and OncoKB. Approximately 20% of the proteins with STARMAP-defined regions exhibit significant separation of survival outcomes, about 10% of mutations have frequently occurred mutations, and approximately 5% of mutations in these top proteins have levels of evidence for clinical or therapeutic utility. **h**, Enrichment analysis of the top 10% variants identified by STARMAP vs. the bottom 90% across clinical annotation databases (STRING, TCGA, ClinVar, OncoKB).

We next evaluated the clinical relevance of these structure-defined regions. Stratification of KRAS variants by their localization within the identified RTIs and RDIs reveals significantly reduced overall survival compared to other KRAS variants^78^ (***Fig. 6f***), indicating that structural context refines prognostic interpretation beyond gene-level mutation status. Extending this analysis proteome-wide, we find that approximately 20% of proteins contain region-defined variants that significantly stratify patient survival^78^, with ∼20% of proteins containing variants that significantly separate clinical outcomes (***Fig. 6g***). STARMAP further identifies additional functionally and clinically relevant regions ***(Extended Data 8)***, and variants within these regions are enriched in curated clinical datasets, including ClinVar^79^ and OncoKB^80,81^. Quantitative analysis shows significant enrichment and elevated odds ratios for these datasets, with strongest signals observed for ClinVar and OncoKB annotations ***(Fig. 6g–h)***. Importantly, these associations are not driven by tissue-specific biases, indicating that STARMAP captures robust mutation–function relationships across contexts ***(Extended Data 9)***.

## DISCUSSION

For more than a century, biochemistry has advanced by uncovering molecular mechanisms through controlled, reductionist experiments^82–88^. These approaches established the structure-function paradigm, revealing how protein architecture governs catalysis,^89–92^ binding,^93–97^, and degradation^98–102^. However, modern biology increasingly confronts systems whose scale and complexity exceed the limits of these approaches^103–107^. Advances in sequencing,^108–111^ imaging^112,113^, and multi-omics^114,115^ technologies now enable the simultaneous measurement of thousands of molecular components and interactions within living cells. While these data provide unprecedented descriptive power, they introduce a central challenge: how can mechanistic insight be extracted when systems can no longer be interrogated one molecule at a time?

A key limitation is not the lack of data, but the lack of mechanistically meaningful representations. Current large-scale analyses identify statistical associations between genes, pathways, function^35,36,116–123,^, but these relationships are often disconnected from the underlying molecular architecture that governs biological processes. Although the principle that three-dimensional molecular organization defines function is well established^21,124–126^, it has not been systematically extended to population-scale datasets, where millions of naturally occurring variants encode rich but unstructured perturbation information^116,124,125^.

STARMAP addresses this gap by establishing protein structure as a coordinate system for interpreting genetic variation. By organizing variants according to spatial proximity, rather than sequence or frequency alone, STARMAP enables systematic identification of functional regions across the proteome, including in previously uncharacterized proteins and variants. This representation reveals how localized perturbations propagate through regulatory networks, enabling the resolution of convergent network behaviors and the identification of molecular states associated with therapeutic sensitivity. More broadly, these results demonstrate that embedding biophysical context into large-scale datasets enables mechanistic insights that are otherwise inaccessible from transcriptomic or genomic data alone.

Recent advances in AI-based models of gene regulation have improved interpretability by incorporating prior biological knowledge, such as transcription factor binding motifs, protein-protein interactions, and gene ontology annotations^53,127–129^. However, most approaches remain limited to gene-level representations and do not account for how distinct regions within a protein uniquely influence regulatory activity. Structural AI models have demonstrated the value of residue-level features for predicting protein function^116,130,131^, but are not designed to infer regulatory relationships or network behavior. STARMAP complements these efforts by introducing a structure-informed representation that enables regulatory and pharmacologic inference directly from protein architecture.

In addition to enabling mechanistic inference, STARMAP provides a scalable strategy for prioritizing functional regions from vast mutational search spaces. Comprehensive mutational and perturbation datasets^16,35,36^ remain available for only a small number of proteins, including BRCA,^16,132^ TP53,^16,133^ and KRAS,^16,134^ reflecting the difficulty of scaling exhaustive experimental approaches. By contrast, STARMAP narrows this search space by identifying regions most strongly associated with regulatory and pharmacologic phenotypes, enabling targeted, hypothesis-driven experimentation. The strong concordance observed across independent datasets, including Perturb-seq^16,17^, single-cell drug perturbation^18^, MAVE,^35,36^ STRINGDB^41–52^, protein-protein interaction databases^53,129^, and clinical cohorts^78–81^, supports the biological relevance of these prioritized regions. Importantly, STARMAP not only identifies candidate regions, but also predicts their likely functional consequences, allowing experiments to be designed with specific mechanistic hypotheses rather than relying on trial-and-error discovery.

Together, these findings extend the structure-function paradigm from individual molecules to complex biological systems, enabling mechanism discovery at scale. By linking protein structure to regulatory networks and cellular phenotypes, STARMAP provides a foundation for a new class of interpretable, structure-informed models that bridge statistical learning and biological mechanisms, opening new opportunities for discovery in systems biology, functional genomics, and precision medicine.

## Data Availability

All scripts to conduct this analysis and reproduce the accompanying figures are available on GitHub (https://github.com/Brunk-Lab/STARMAP_paper). The STARMAP pipeline is also available as a downloadable software package on Github (https://github.com/Brunk-Lab/STARMAP). An interactive web resource accompanying this work enables direct exploration of structure-resolved associations across proteins, regulatory networks, and drug responses, providing a platform for hypothesis generation, functional interpretation, and discovery (available at: starmap.unc.edu).

## METHODS

### Data sources and integration

Genomic variation and transcriptomic data were obtained from DepMap/CCLE^10–12^ (v24Q4), which includes somatic variants and RNA-seq profiles across ∼1,700 cancer cell lines. Single-cell perturbation datasets, including loss-of-function^17^ and knock-in Perturb-seq experiments^16^, were integrated to enable independent validation of TRN activity. Drug response data were obtained from the CTRP^13^, and additional single-cell drug perturbation profiles were sourced from the Tahoe-100M dataset^18^.^35,36^ data^35,36^ were used to assess functional impact at the protein level. All datasets were aligned at the cell line level to ensure matched genotype–phenotype comparisons across modalities.

### Construction of STARMAP representations

Variants were mapped onto protein structures derived from the Protein Data Bank^15^ and AlphaFold^14^ models. For each variant, residue-level coordinates corresponding to the Cα atom were extracted from structural models. High-confidence mappings were retained by filtering for sequence similarity, alignment quality, and structural confidence (pLDDT ≥ 70 for AlphaFold). Variants located in disordered or poorly resolved regions were excluded. This resulted in a structurally resolved dataset of variants across thousands of proteins, forming the basis for downstream spatial analyses.

For each protein, residue-level 3D coordinates were obtained from a precompiled structure table (derived from PDB^15^ or AlphaFold^14^ models) containing the Cartesian coordinates of the alpha Carbon atom for each residue. Residues were used as the fundamental units for downstream embedding and clustering.

To reduce sensitivity to absolute coordinate scale and to ensure comparability across proteins, residue-level structural features were rank-normalized prior to dimensionality reduction. Briefly, coordinate values were converted to ranks, ranks were mapped to their mean values, and each feature was rescaled to a 0-1 range using min-max normalization. Features with no variance across residues were excluded.

To generate a low-dimensional representation that preserves structural relationships between residues, non-negative matrix factorization (NMF)^135^ was applied to the normalized residue-by-feature matrix.

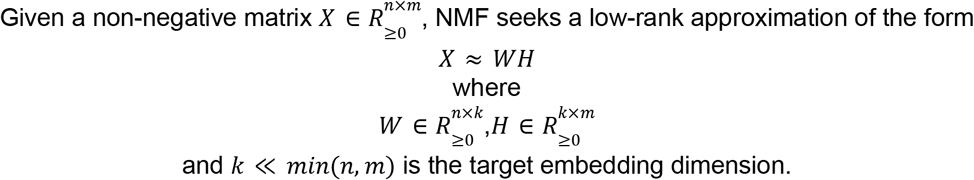

NMF decomposes the structural representation into a set of non-negative latent components that capture dominant spatial patterns within the folded protein. The resulting residue-level latent matrix provides an interpretable embedding in which residues sharing similar three-dimensional environments have similar coordinates in latent space.

The dimensionality of the latent representation was selected independently for each protein. NMF models were fit across a range of candidate component numbers (k>3), and reconstruction error was evaluated using mean squared error between the original and reconstructed structural matrices. The value of k minimizing reconstruction error was selected for the final embedding.

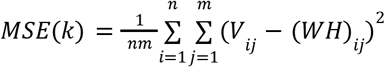

To produce a two-dimensional spatial layout suitable for visualization and clustering, pairwise Euclidean distances between residues were computed in the NMF-derived latent space. These distances were then embedded into two dimensions using metric multidimensional scaling (MDS)^136^, yielding a continuous two-dimensional “flatmap” in which distances between residues reflect similarities in their underlying three-dimensional structural context. The resulting coordinates were rescaled to a common range for consistency across proteins.

An additional residue-level feature, termed altitude, was computed from the latent components to capture deviations from the average structural profile of the protein and was carried forward for visualization and downstream analyses.

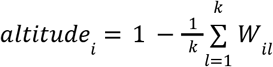

Residues were grouped into spatially coherent regions using k-means clustering^137^ applied to the two-dimensional flatmap coordinates. The number of clusters was set equal to the selected latent dimensionality k, ensuring that region definitions reflect the dominant structural modes of the protein. This procedure partitions each protein into contiguous regions that correspond to shared structural environments rather than linear sequence proximity.

The final flatmap representation consists of two-dimensional coordinates, an altitude value, and a cluster assignment for each residue.Protein-protein interactions from the Integrated Interactions Database^129^ and BioGRID^53^ and annotation data from Uniprot^138^ were used for functional annotation of structural regions.

### Association of structural regions with TRN transcriptional activity

TRNs were identified using >500 transcription factor target^25,26^ sets^25,26^. The gene set for each transcription factor was considered its TRN. TRN transcriptional activity was quantified using single-sample gene set enrichmen^23,24^sis^23,24^ across these gene sets. For each protein, variant positions were aligned with TRN activity measurements across matched cell lines. Spatial association between variant-localized signals and TRN activity was quantified using the Getis–Ord Gi* statistic^22^, which measures local enrichment relative to the global distribution .

For each protein, two-dimensional residue coordinates and cluster assignments were obtained from the corresponding structural flatmap. Transcriptional regulatory network (TRN) activity scores were derived from gene set enrichment analyses (GSEA) of cancer cell line transcriptomes.

For a given protein-TRN pair, mutation positions observed across cell lines were matched to TRN activity scores associated with those same cell lines. Each residue position was thus assigned one or more TRN scores corresponding to independent mutation events at that residue. Residues without observed mutations for a given TRN were assigned a score of zero. When multiple TRN scores were associated with a single residue, scores were retained explicitly rather than averaged. To accommodate this, residue-level TRN scores were expanded into multiple aligned columns, each representing one mutation-associated score. Missing values were filled with zero, yielding a sparse residue-by-score matrix defined on the protein flatmap.

Residue coordinates from the flatmap were represented as a planar point set for further analysis. Spatial relationships between residues were defined using Queen contiguity weights^139^, which consider residues to be neighbors if they share a local adjacency in the two-dimensional embedding. This neighborhood structure defines the spatial window over which local clustering of TRN-associated signal is assessed.

Spatial association between TRN activity and protein structure was quantified using the Getis-Ord local statistic (Gi*)^22^. For each residue i and each TRN score column j, the Gi* statistic was computed as a standardized z-score:

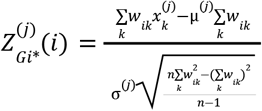

Where 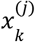 is the TRN score at residue k for column j, *w*_*ik*_ denotes the spatial weight between residues i and k, and µ ^(*j*)^ and σ_(*j*)_ are the mean and standard deviation of the TRN scores across all residues for that column

This formulation produces a standardized measure of local enrichment or depletion relative to the global distribution of scores on the protein flatmap.

To obtain a single residue-level statistic per TRN, Gi* z-scores were summed across all mutation-associated columns:

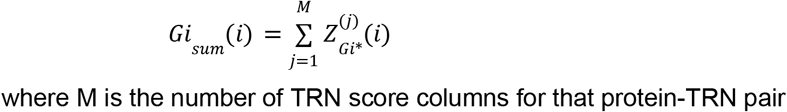

This aggregation captures the cumulative spatial signal arising from multiple independent mutation events while preserving the standardized nature of the underlying statistic.

Residue-level Gi-sum values were mapped back to the structure-derived clusters defined on the flatmap. For each cluster c, a cluster-level TRN score was computed as the mean of the residue-level values:

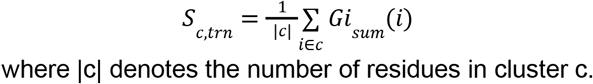

In parallel, the number of residues within each cluster exhibiting positive local signal Gi-sum>0 was recorded as a coverage metric reflecting the spatial extent of TRN-associated enrichment within the region.

Several normalization steps ensure that TRN Gi* scores are comparable across proteins of different sizes, topologies, and mutation burdens. First, the use of z-scored Gi* statistics standardizes local enrichment relative to each protein’s own spatial and score distributions. Second, cluster-level scores are computed as means rather than sums, preventing larger clusters from dominating purely due to residue count. Third, residues without mutation-associated TRN scores are explicitly assigned zero prior to Gi* computation, establishing a consistent baseline across proteins. Together, these steps enable direct comparison of spatial TRN enrichment patterns across proteins within the STARMAP framework.

Only high-confidence associations (top 10% of Gi* scores) were retained for downstream analyses.

### Association of structural regions with drug response

Drug sensitivity was modeled using a structure-aware machine learning framework. For each variant, spatial features were derived, including cluster identity and distances to annotated functional sites obtained from Uniprot^138^. Drug response labels were defined using CTRP^13^ AUC values, binarized based on percentile thresholds. A multilayer perception-based deep learning model^54^ was trained to predict drug sensitivity from variant-level features using group-aware cross-validation to prevent leakage across cell lines.

To identify drugs that preferentially associate with specific regions of a protein, a cluster-level enrichment analysis was performed using the predefined clusters from flatmaps and the predictions from the multilayer perceptron.

Because individual residues may appear multiple times in the prediction outputs, predictions were first aggregated at the residue level by averaging predicted probabilities across replicates. Residues were classified as “high-confidence” for a given drug if their mean predicted probability exceeded a fixed threshold of 0.90:

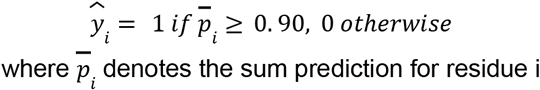

For each protein–drug pair, enrichment of high-confidence residues within a given cluster k was evaluated relative to the rest of the protein. A 2×2 contingency table was constructed by counting residues inside and outside cluster k that were classified as high-or low-confidence:

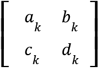

where *a*_*k*_ and *b*_*k*_ are the numbers of high- and low-confidence residues within cluster k, respectively, and *c*_*k*_ and *d*_*k*_ are the corresponding counts for all other residues in the same protein

Fisher’s exact test was applied to this table to quantify whether cluster k was enriched for drug-associated residues compared with the remainder of the protein. The strength of enrichment was summarized using the log-transformed odds ratio:

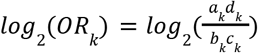

In cases where zero counts led to infinite odds ratios, the corresponding log2 odds ratio was capped at an absolute value of 6 to prevent domination by singular estimates.

### Network and global analyses

Protein-TRN and protein–drug associations were represented as bipartite networks and projected into protein–protein similarity graphs based on shared associations. Network metrics, including degree, centrality, and clustering, were used to characterize regulatory breadth and amplitude. Similarity between TRNs and drugs with high TRN association scores (GI*) and high drug sensitivity association scores (log-odds) was computed using Jaccard indices, enabling identification of regulatory and pharmacologic modules. Functional enrichment analyses were performed using Gene Ontology^127,128^ annotations to interpret network structure.

### Validation and benchmarking

STARMAP predictions were validated using independent Perturb-seq datasets^16,17^ and single-cell drug response datasets^18^ by comparing predicted TRN rankings with experimentally observed transcriptional responses. Agreement was quantified using rank-based confidence metrics and classification performance measures (AUROC, AUPRC, F1 score, and accuracy). Additional validation was performed ^35,36^ MAVE^35,36^ data and PolyPhen^6^ annotations to assess functional impact at the protein level. STARMAP performance was benchmarked against established gene regulatory network^37–39^ and drug response prediction^55–57^ methods. Clinical relevance was assessed by comparing STARMAP-defined regions with patient survival data^78^ and curated variant databases, including ClinVar^79^ and OncoKB^80,81^.

## Acknowledgements

The authors thank Jeff Roach and the UNC Research Computing team for their support in pipeline development and computing resources, and Carolina CloudApps for hosting the accompanying interactive web resource. The authors also acknowledge Pablo Tamayo for the many helpful discussions that made this work possible. K.S. and J.C. were supported by funding from the PhRMA Foundation and the Lung Cancer Research Foundation Boehringer Ingelheim Early Career Investigator Award.

## Author Contributions

Conceptualization, E.B. ; methodology, E.B., K.S., J.C. ; formal Analysis, K.S., J.C., E.B.; funding Acquisition, E.B.; investigation, K.S., J.C., E.B.; resources, E.B.; supervision, E.B.; validation, K.S., J.C.; visualization, K.S., J.C., E.B.; writing-original draft, E.B., K.S.; writing-review and editing, All authors. All authors have read and agreed to the published version of the manuscript.

## Inclusion and Ethics Statement

All experiments involving cell lines were conducted in compliance with institutional biosafety and research integrity guidelines approved by the University of North Carolina at Chapel Hill. No human or animal subjects were involved in this study. We are committed to fostering a supportive research environment. Our team reflects a range of backgrounds and training levels, and we actively mentor junior scientists across disciplines. We strive to ensure that our research practices are transparent, reproducible, and accessible to the broader scientific community. Data and code from this work are openly shared to promote collaboration and accountability.

## Extended Data Figures

**Extended Data 1:**
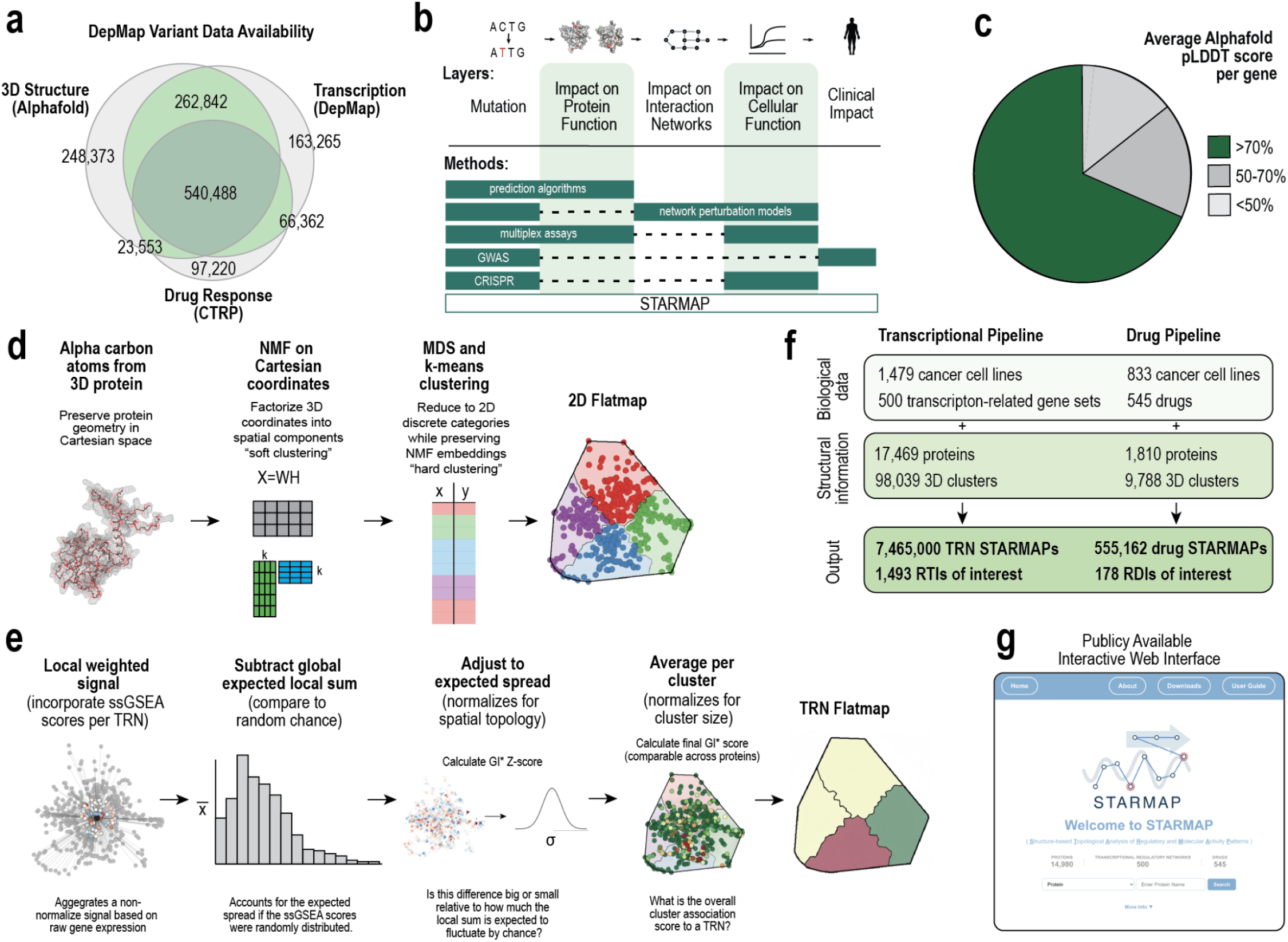
Overview of the STARMAP framework, data integration, and analytical workflow. **a**, Overlap of data modalities used in this study across DepMap resources, including 3D protein structures (AlphaFold), transcriptional profiles (DepMap), and drug response data (CTRP). The Venn diagram indicates the number of variants supported by each modality and their intersections. **b**, Conceptual schematic illustrating the propagation of somatic mutations across biological scales, from protein function to interaction networks, cellular phenotypes, and ultimately clinical impact. Existing experimental and computational approaches (e.g., prediction algorithms, multiplex assays, genome-wide association studies, CRISPR, and network perturbation models) are shown in relation to these layers, highlighting the integrative scope of STARMAP. **c**, Distribution of average AlphaFold pLDDT confidence scores per protein, demonstrating that the majority of proteins analyzed have high-confidence structural predictions (>70%). **d**, Structural embedding and clustering workflow. Alpha carbon coordinates from 3D protein structures are decomposed using non-negative matrix factorization to obtain spatial components, followed by dimensionality reduction and *k*-means clustering to define discrete regions of functional interest on a 2D flatmap representation. **e**, Statistical framework for identifying TRN associations. Local weighted signals derived from ssGSEA scores are compared to a null distribution, adjusted for spatial topology using a Getis–Ord Gi*\** statistic, and aggregated at the cluster level to generate normalized association scores comparable across proteins. **f**, Scale of analyses performed. The transcriptional pipeline integrates 1,479 cancer cell lines and ∼500 TRN gene sets across 17,469 proteins, yielding ∼7.5 million TRN-specific STARMAPs. The drug pipeline includes 833 cell lines and 545 drugs across 1,810 proteins, producing over 550,000 drug-specific STARMAPs. Outputs include prioritized RTIs and RDIs. **g**, Publicly accessible STARMAP web interface, enabling exploration of protein structural maps, TRN associations, and drug response signatures.

**Extended Data 2:**
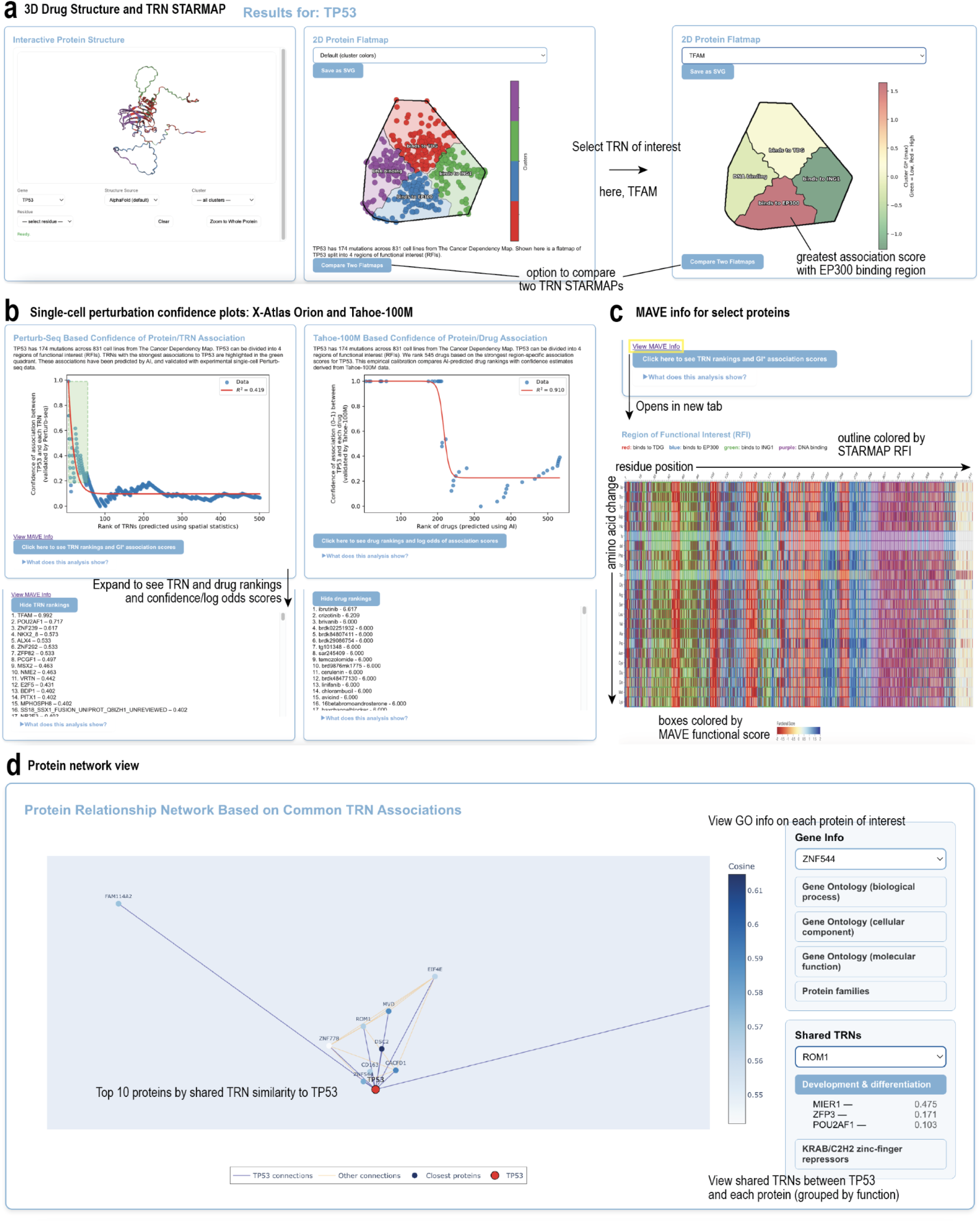
Extended Data STARMAP web interface for interactive exploration of protein structure, transcriptional programs, and drug associations. **a**, Interactive visualization of STARMAP outputs for a representative protein (TP53). The interface integrates 3D protein structure, 2D flatmap clustering, and TRN-specific STARMAP overlays, allowing users to select TRNs (e.g., TFAM) and identify regions of functional interest with the strongest association signals. Users can also compare multiple TRNs. **b**, Single-cell perturbation–based confidence plots derived from Perturb-seq (X-Atlas Orion) and Tahoe-100M datasets. These plots show the relationship between STARMAP-predicted TRN rankings and experimental confidence, with expandable panels providing detailed TRN and drug rankings and confidence and log-odds scores. **c**, Integration of MAVE data for select proteins. Residue-level functional scores are visualized for STARMAP-defined regions of functional interest, enabling comparison between experimental mutational effects and predicted transcriptional associations. **d**, Protein–protein network view based on shared TRN associations. The interface displays nearest neighbors (e.g., top proteins most similar to TP53) using cosine similarity, with options to explore Gene Ontology annotations, protein families, and shared TRNs grouped by functional categories.

**Extended Data 3:**
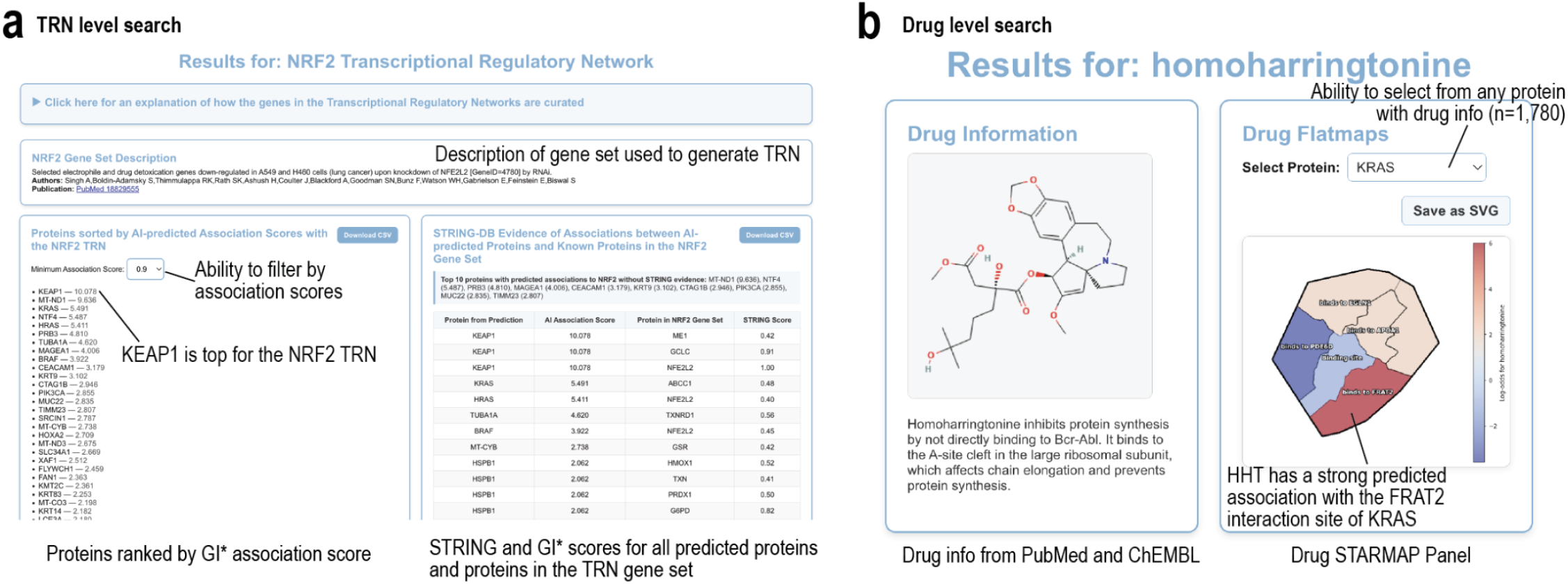
Extended Data STARMAP web interface for transcriptional network and drug-centric exploration. **a**, TRN-level search interface illustrated for the NRF2 TRN. The platform provides curated gene set descriptions and proteins ranked based on STARMAP Gi*\** association scores with filtering by minimum score thresholds. Integrated evidence from STRING highlights known and predicted interactions between top-ranked proteins and the TRN gene set, enabling prioritization of key regulators (e.g., KEAP1). **b**, Drug-level search interface illustrated for homoharringtonine. The platform integrates chemical structure and functional annotations with protein-specific drug STARMAPs, allowing users to select proteins of interest and visualize spatial drug–protein associations. In this example, homoharringtonine is predicted to have a strong association with a KRAS interaction region, demonstrating the utility of STARMAP for linking drugs to structural regions of functional relevance.

**Extended Data 4:**
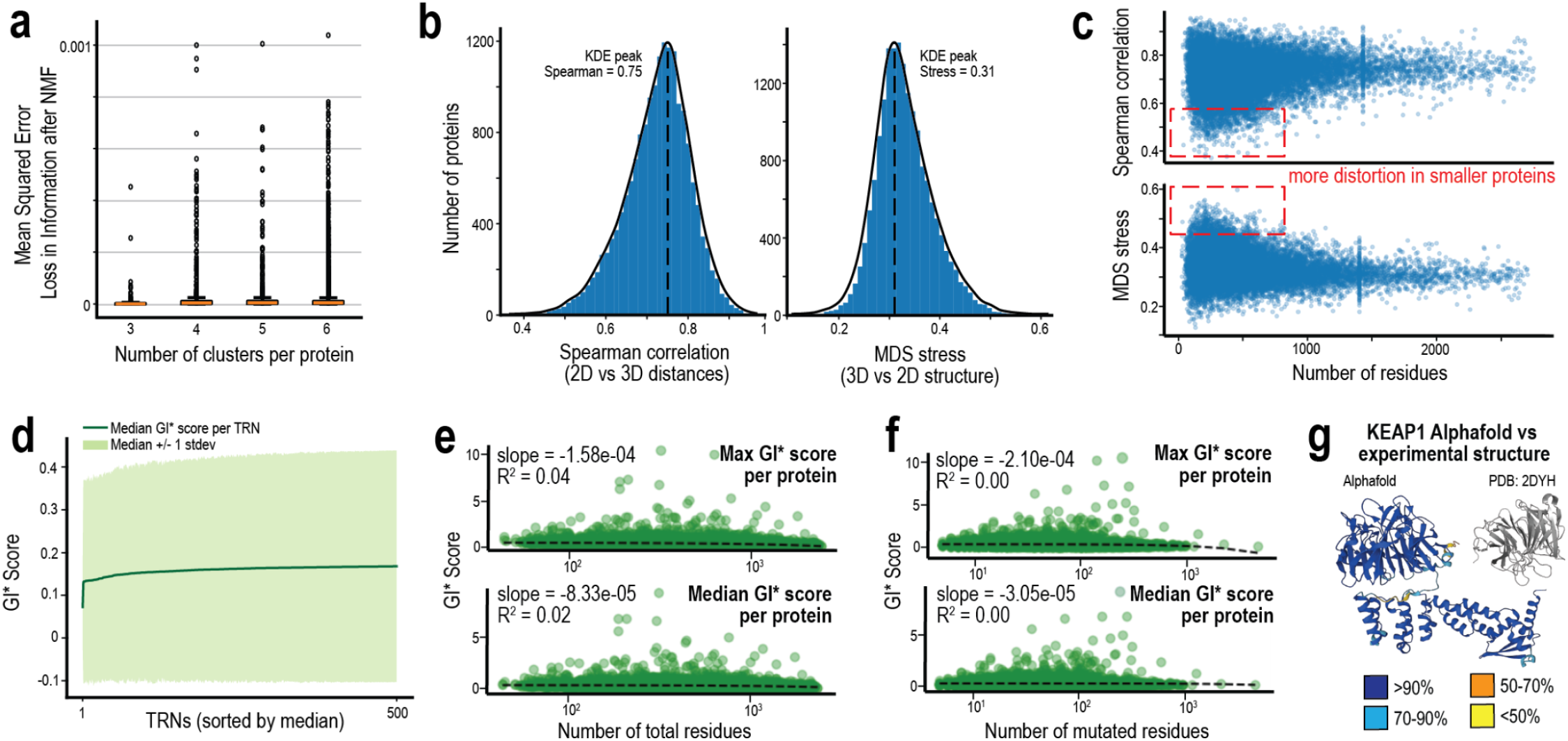
Extended Data Validation of structural embeddings and robustness of STARMAP-derived signals. **a**, Reconstruction error following non-negative matrix factorization of protein structures as a function of the number of clusters (*k* = 3–6). Mean squared error is low across cluster choices, indicating minimal information loss during dimensionality reduction. **b**, Fidelity of 2D structural embeddings relative to original 3D geometry. Left: distribution of Spearman correlations between pairwise residue distances in 2D versus 3D space (median ≈ 0.75), demonstrating strong preservation of spatial relationships. Right: Distribution of multidimensional scaling (MDS) stress values (median ≈ 0.31), indicating low embedding distortion. **c**, Relationship between protein size (number of residues) and embedding quality. Smaller proteins have greater variability and distortion, reflected by lower Spearman correlations and higher MDS stress, whereas larger proteins show more stable embeddings. **d**, Distribution of normalized TRN association scores (Gi*) across TRNs, sorted by median score. Shaded regions represent ±1 standard deviation, demonstrating consistent scaling and comparable signal distributions across TRNs. **e**, Dependence of TRN-associated signal strength on protein size. Maximum and median Gi*\** scores per protein show minimal correlation with total residue count (low slopes and R2), indicating that STARMAP signals are not driven by protein length. **f**, Dependence of TRN-associated signal strength on mutation burden. Maximum and median Gi*\** scores per protein show negligible association with the number of mutated residues, supporting robustness to variation in mutation frequency. **g**, Structural consistency between AlphaFold predictions and experimentally resolved structures, illustrated for KEAP1 (PDB: 2DYH). High-confidence regions (pLDDT >90%) align closely with the experimental structure, supporting the reliability of AlphaFold-derived inputs for downstream analyses.

**Extended Data 5:**
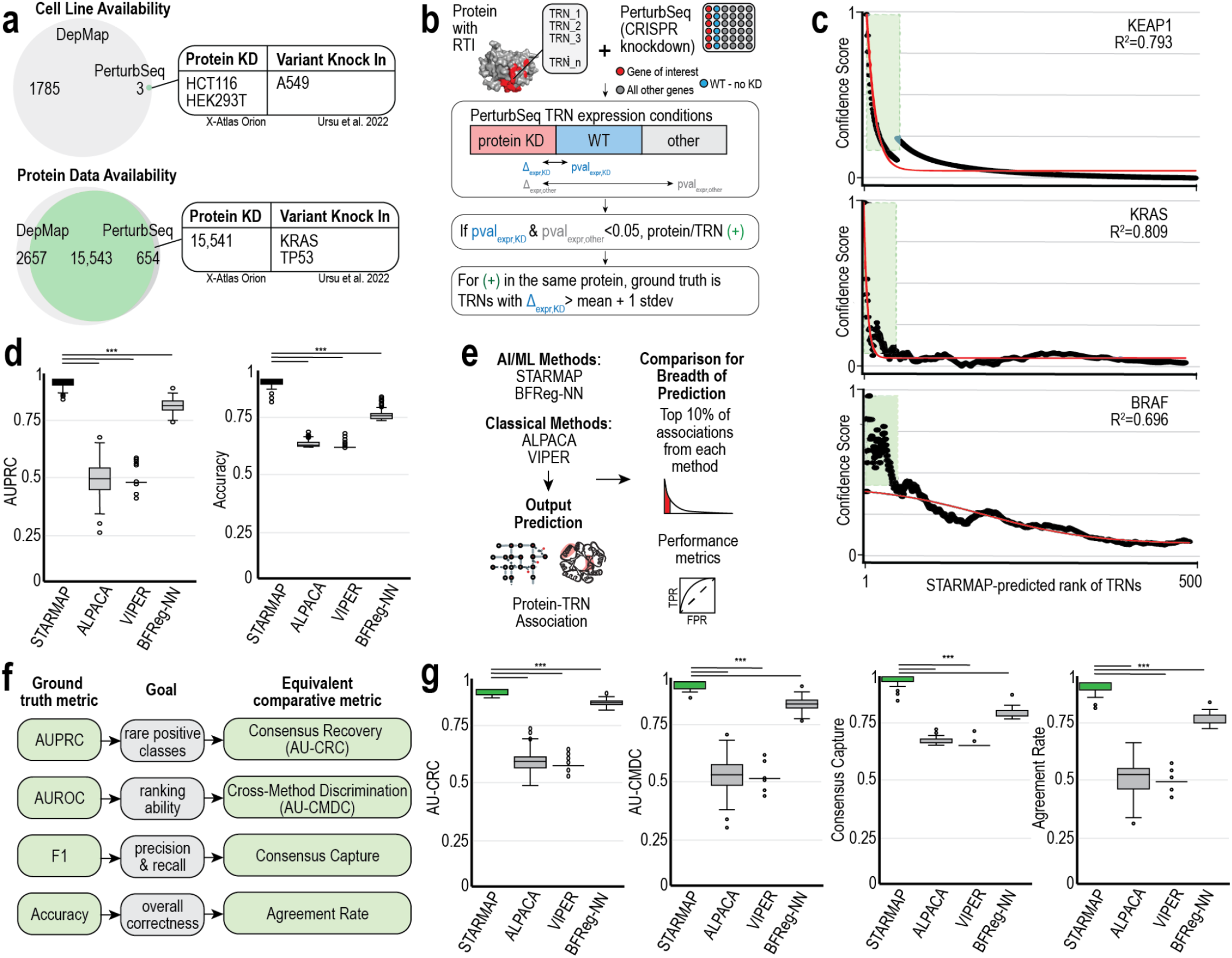
Extended Data Benchmarking framework and validation of STARMAP predictions using Perturb-seq and comparative methods. **a**, Availability of matched cell line and protein-level perturbation data across DepMap and Perturb-seq resources. Overlap in cell lines and proteins enables benchmarking using CRISPR-based perturbation datasets, including protein knockdown (KD) and variant knock-in experiments. **b**, Strategy for defining ground truth TRN associations using Perturb-seq. For proteins with RTIs, TRN transcriptional activity is compared across protein knockdown, wild-type (WT), and other conditions. TRNs are labeled as positive if their differential expression relative to both WT and background conditions is significant (p < 0.05). Ground truth positives are further refined within each protein as TRNs with expression shifts exceeding one standard deviation. **c**, Empirical validation of STARMAP confidence scores using Perturb-seq data. For representative proteins (KEAP1, KRAS, BRAF), STARMAP-predicted TRN rankings show strong concordance with experimentally derived confidence scores, with high goodness-of-fit (R^2^ values). The enrichment of true positives among top-ranked TRNs (highlighted region) demonstrates effective prioritization. **d**, Comparison of STARMAP with existing methods (ALPACA, VIPER, BFReg-NN) using area under the precision–recall curve (AUPRC) and accuracy. STARMAP consistently achieves higher predictive performance, particularly in identifying true positive TRN associations. **e**, Overview of benchmarking design. AI/machine learning-based methods (STARMAP, BFReg-NN) and classical network-based approaches (ALPACA, VIPER) were evaluated based on their abilities to predict protein–TRN associations. Comparisons focused on the top 10% of predicted associations per method, emphasizing breadth and prioritization performance. **f**, Ground-truth metric, goals, and comparative metric. **g**, Comparative performance across consensus-based metrics.

**Extended Data 6:**
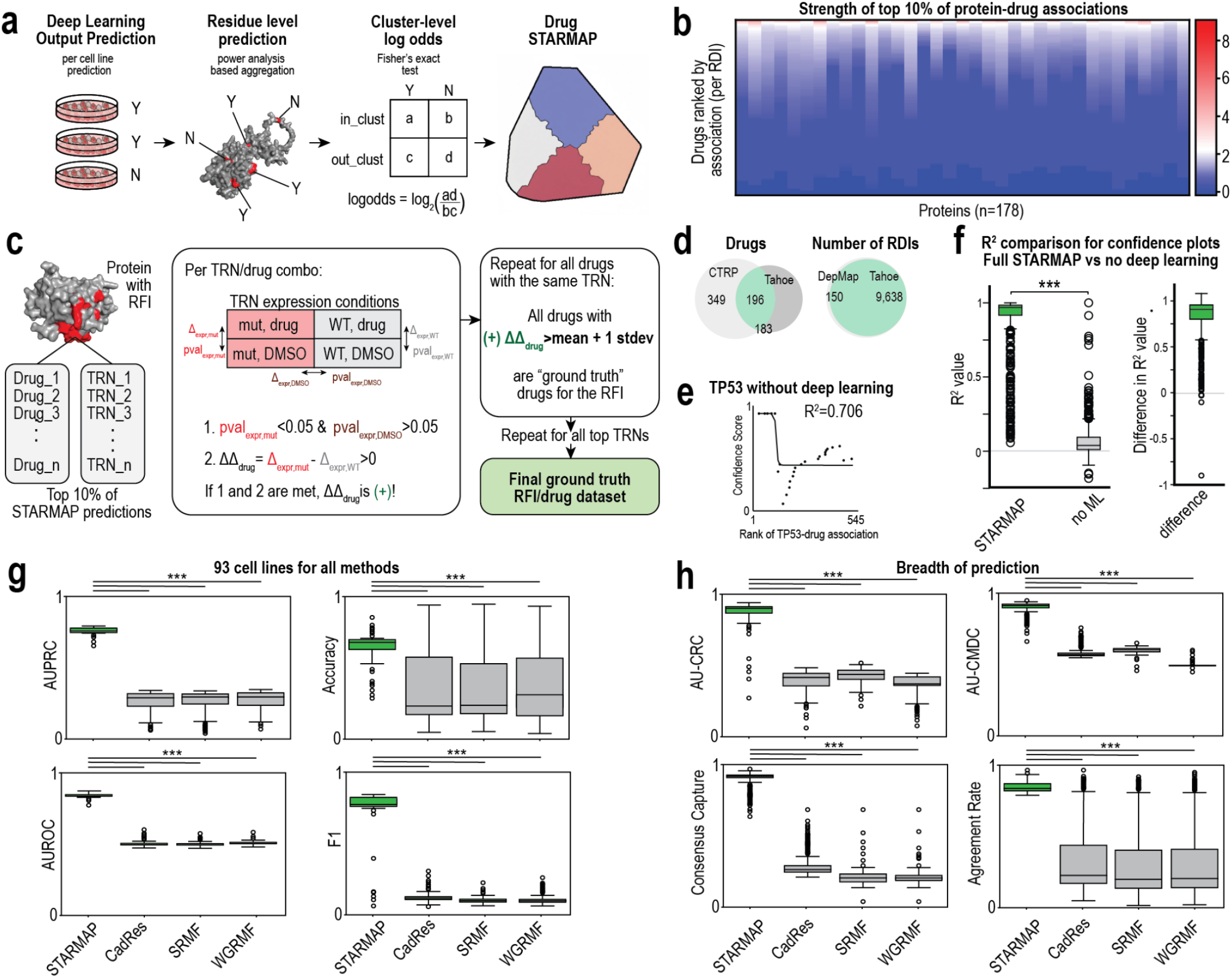
Extended Data Drug-level STARMAP framework, validation strategy, and benchmarking of predictive performance. **a**, Workflow for generating drug-specific STARMAPs. Deep learning–based predictions were first computed at the cell line level and aggregated to residue-level signals. These were summarized at the cluster level using Fisher’s exact test to compute log-odds ratios, which were then mapped onto 2D structural flatmaps to produce drug-specific STARMAPs. **b**, Heatmap showing the strength of the top 10% of protein–drug associations across proteins (n = 178), with drugs ranked per RDI. Color intensity reflects log-odds–based association strength. **c**, Strategy for defining ground truth drug–TRN associations using Perturb-seq. For each TRN–drug pair, differential expression was evaluated across mutant and wild-type conditions under both drug treatment and DMSO controls. Associations were considered positive if (i) drug-specific perturbation was significant (p < 0.05) and DMSO treatment was not and (ii) the differential effect exceeded that of wild-type. Ground truth drugs for a given RDI were defined as those with effects exceeding one standard deviation above the mean across all drugs. **d**, Left: intersection between CTRP and Tahoe drug sets. Right: Number of RDIs identified using DepMap versus Tahoe datasets, highlighting expanded coverage in Tahoe. **e**, Example validation of TP53 without deep learning integration, showing reduced concordance between predicted rankings and empirical confidence scores (lower R^2^). **f**, Comparison of confidence calibration performance with and without deep learning. Full STARMAP yields significantly higher R^2^ values relative to models without machine learning, demonstrating improved alignment with experimental data. **g**, Benchmarking across 93 shared cell lines comparing STARMAP to alternative approaches (CadRes, SRMF, WGRMF). STARMAP achieves superior performance across AUPRC, accuracy, AUROC, and F1 score. **h**, Evaluation of prediction breadth and cross-method consistency. STARMAP outperforms other methods in consensus-based metrics, including consensus recovery (AU-CRC), cross-method discrimination (AU-CMDC), consensus capture, and agreement rate, indicating stronger prioritization and agreement across independent approaches.

**Extended Data 7:**
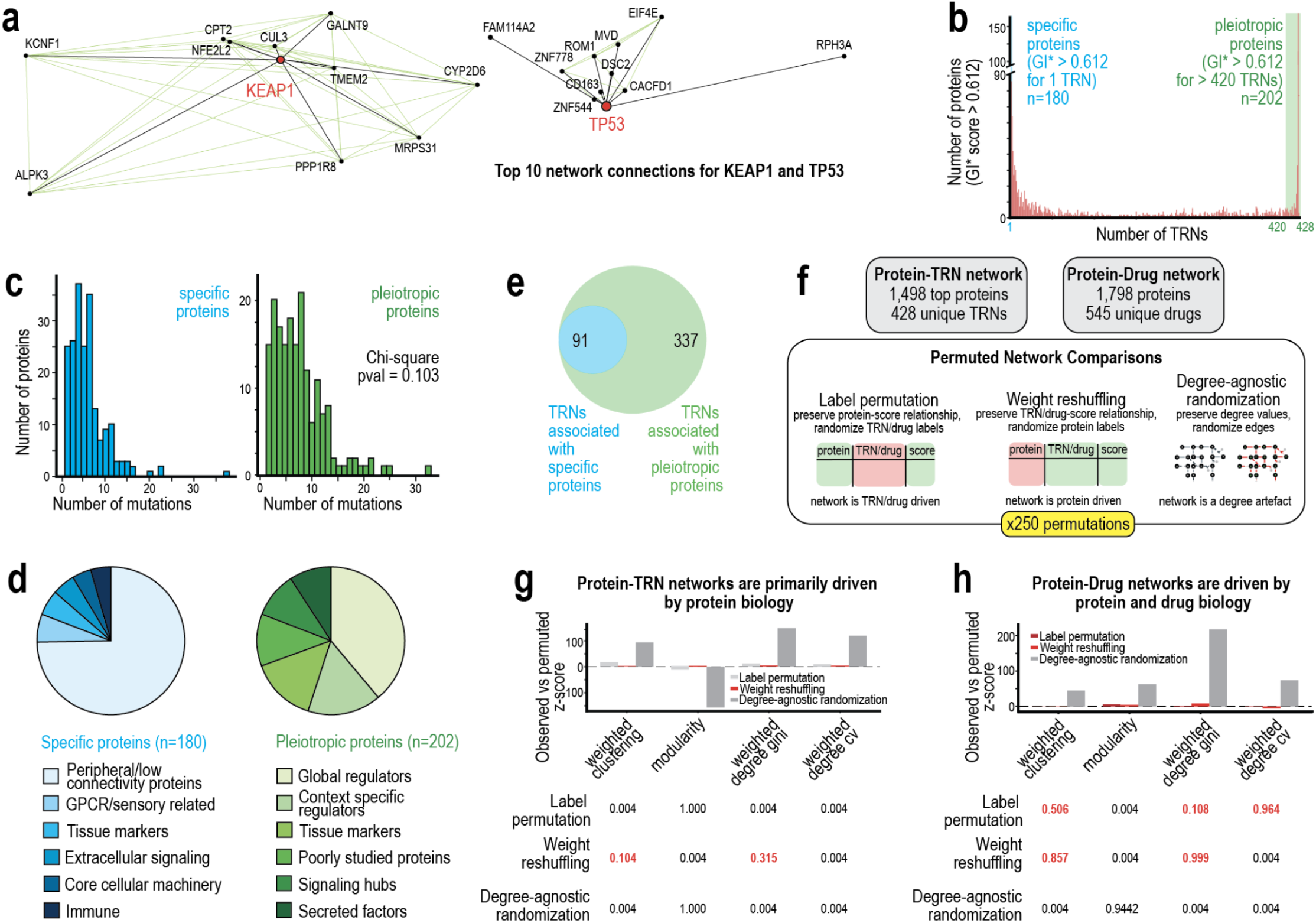
Extended Data Network structure, protein specificity, and drivers of STARMAP-derived protein-TRN and protein-drug associations. **a**, Top network connections for representative proteins KEAP1 and TP53 within the protein-TRN interaction network. Nodes represent proteins and edges reflect shared or correlated TRN associations, highlighting distinct connectivity patterns between proteins. **b**, Distribution of the number of TRNs associated with each protein (defined by TRN association score - Gi*>0.612). Proteins segregate into “specific” proteins with associations to a single TRN (n = 180) and “pleiotropic” proteins associated with a large number of TRNs (>420; n = 202), indicating heterogeneity in regulatory breadth. **c**, Histograms of the distribution of mutation counts for specific versus pleiotropic proteins; no significant difference was observed (chi-square p = 0.103), suggesting that pleiotropy is not driven by mutation frequency. **d**, Functional annotation of protein classes. Specific proteins are enriched for peripheral or specialized roles (e.g., GPCR/sensory, tissue markers), whereas pleiotropic proteins are more frequently global regulators, signaling hubs, or broadly acting factors. **e**, Overlap of TRNs associated with specific versus pleiotropic proteins. A subset of TRNs are shared (n = 91), but most TRNs are uniquely associated with pleiotropic proteins (n = 337). **f**, Network construction and permutation framework. Protein-TRN (1,493 proteins, 428 TRNs) and protein–drug (1,798 proteins, 545 drugs) networks were analyzed using three null models: label permutation (preserving protein–score relationships), weight reshuffling (preserving TRN/drug–score relationships), and degree-agnostic randomization. Each comparison was repeated across 250 permutations. **g**, Permutation analysis of protein–TRN networks. Observed network statistics (e.g., clustering, modularity, degree distribution) indicate that network structure is primarily driven by intrinsic protein biology, as label permutation significantly disrupted structure and other null models had limited impact. **h**, Permutation analysis of protein-drug networks. Both protein- and drug-level factors contribute to network structure, as evidenced by sensitivity to label permutation and weight reshuffling, and the minimal effect of degree-preserving randomization. Thus, observed structure is not a trivial artifact of network topology.

**Extended Data 8:**
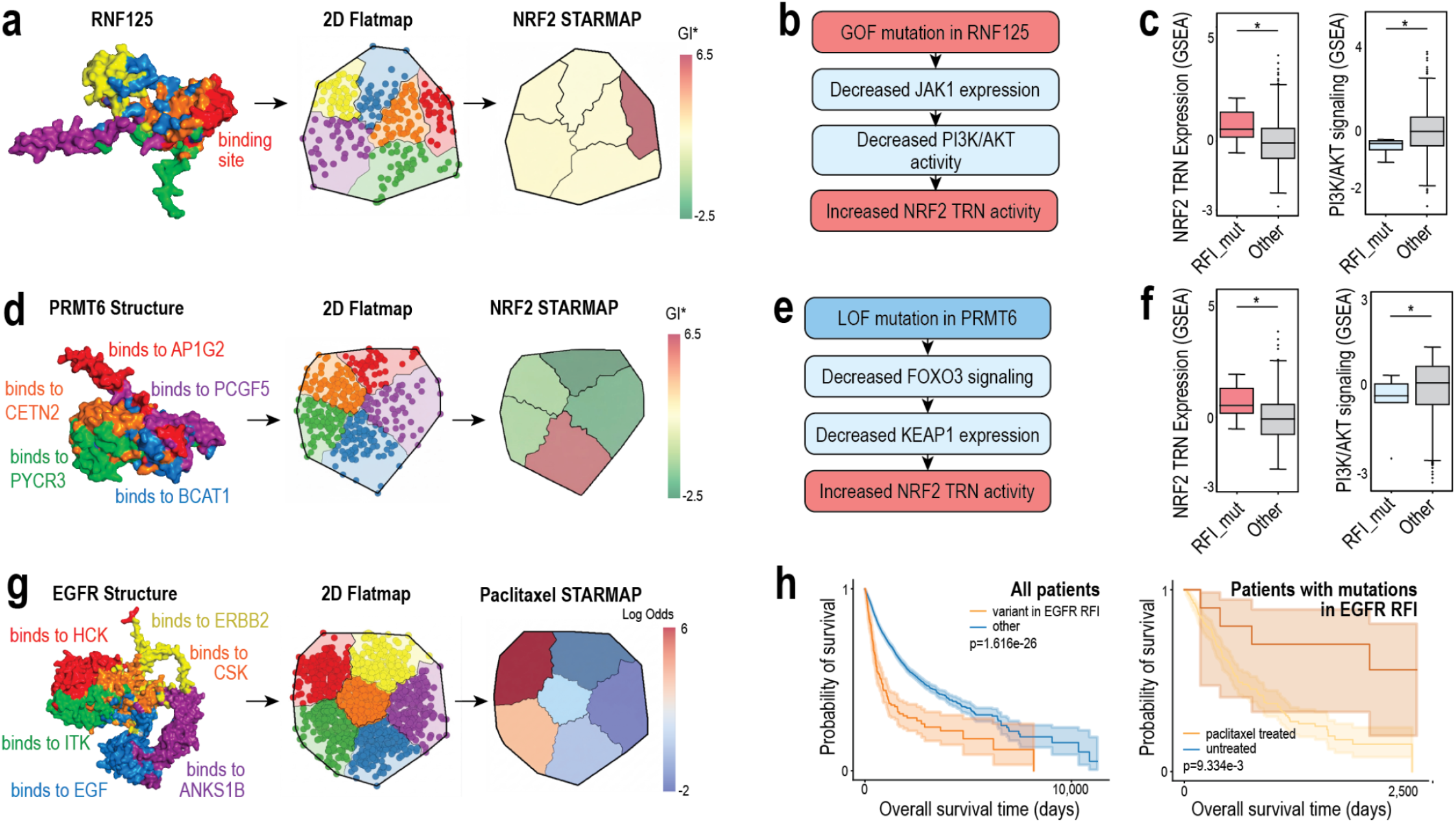
Extended Data Mechanistic interpretation and clinical validation of additional STARMAP-derived protein-TRN and protein-drug associations. **a**, Structural mapping and STARMAP analysis for RNF125. Residue-level clustering was projected onto a 2D flatmap and used to compute NRF2 TRN association scores - Gi*, highlighting an RTI near a predicted binding interface. **b**, Proposed mechanism linking RNF125 gain-of-function mutations to NRF2 activation. RNF125 mutations are associated with decreased JAK1 expression and reduced PI3K/AKT signaling, culminating in increased NRF2 TRN activity. **c**, Validation of RNF125-associated effects using gene set enrichment analysis (GSEA) on transcriptomics data from DepMap. Mutations within the RTI are associated with increased NRF2 TRN expression and decreased PI3K/AKT signaling relative to other mutations. **d**, Structural mapping and STARMAP analysis for PRMT6. Distinct structural clusters correspond to known binding interfaces with multiple interaction partners, with STARMAP identifying regions linked to NRF2 TRN modulation. **e**, Proposed mechanism for PRMT6 loss-of-function mutations. PRMT6 disruption is associated with decreased FOXO3 signaling and reduced KEAP1 expression, leading to increased NRF2 TRN activity. **f**, Transcriptomics and GSEA validation of PRMT6-associated effects. Mutations within the RTI show increased NRF2 TRN expression and decreased PI3K/AKT signaling compared to other mutations. **g**, Structural and drug-response mapping for EGFR. STARMAP identified spatial regions associated with paclitaxel sensitivity, with clusters corresponding to known binding interactions and signaling interfaces. **h**, Left: Across all patients, mutations within the EGFR RTI are associated with significantly worse overall survival. Right: Among patients with mutations in the EGFR RTI, paclitaxel treatment is associated with improved survival compared to untreated patients, supporting the predictive value of STARMAP for therapeutic stratification.

**Extended Data 9:**
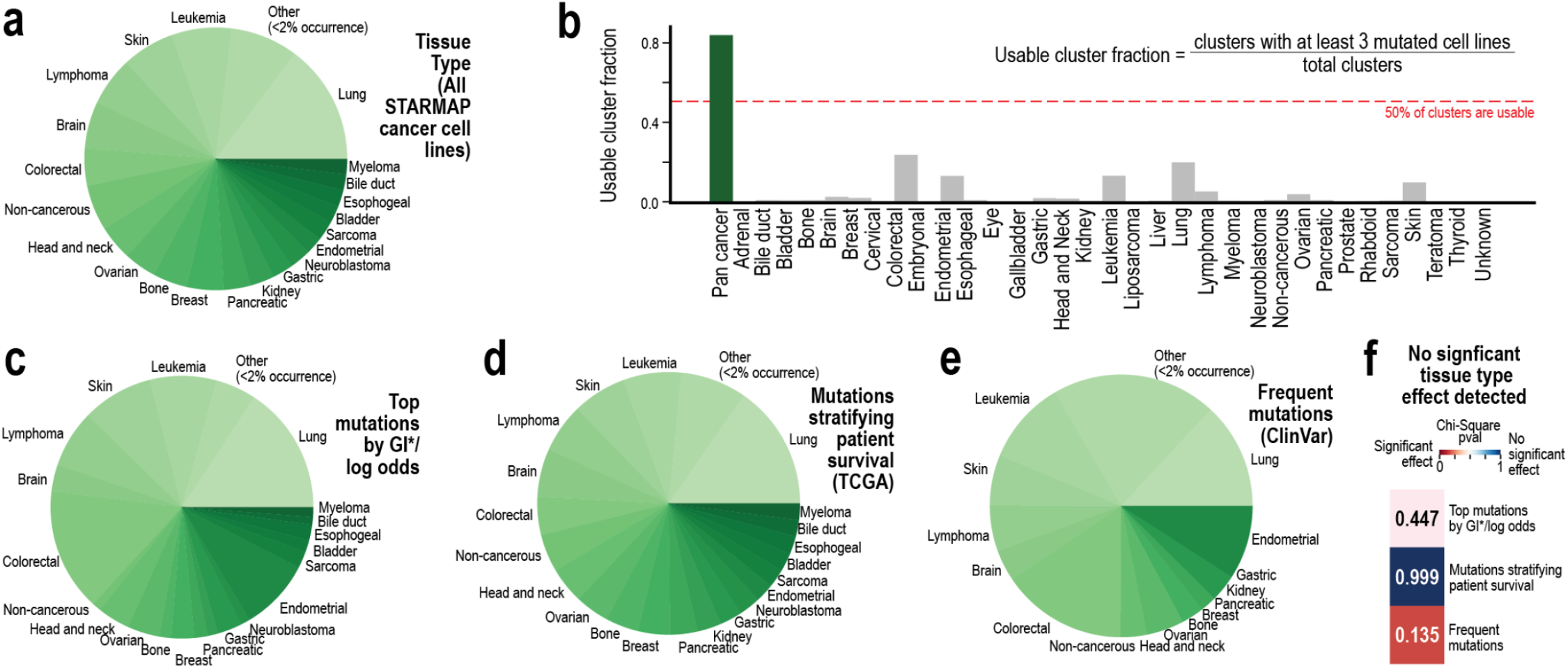
Extended Data Tissue distribution and analysis of tissue type effects in STARMAP’s clinical utility. **a**, Distribution of tissue types represented in the full STARMAP cancer cell line dataset, showing broad coverage across diverse cancer indications. **b**, Fraction of usable clusters per tissue type, defined as clusters with at least three mutated cell lines relative to total clusters. A reference threshold of 50% is indicated. Most tissue types show low fractions, highlighting the need for a pan-cancer approach. **c**, Tissue distribution of top mutations identified by STARMAP (based on TRN association score - Gi*\** and drug sensitivity association score - log-odds), demonstrating similar representation across cancer types as the full dataset. **d**, Tissue distribution of mutations used to stratify patient survival in TCGA, showing comparable coverage across indications. **e**, Tissue distribution of frequently observed mutations from ClinVar, providing an external reference for mutation prevalence across cancer types. **f**. Statistical assessment of tissue-type bias. Chi-square tests indicate no significant enrichment or depletion of tissue types across (i) top STARMAP mutations, (ii) mutations used for survival stratification, or (iii) frequently observed mutations (all p > 0.05), indicating that STARMAP-derived signals are not driven by tissue composition.

## References

1. Zhuang, Y., Wu, F., Chen, C. & Pan, Y. Challenges and opportunities: from big data to knowledge in AI 2.0. Front. Inf. Technol. Electron. Eng. 18, 3–14 (2017).

2. Bruggeman, F. J. & Westerhoff, H. V. The nature of systems biology. Trends Microbiol. 15, 45–50 (2007).

3. Nicholson, D. J. The concept of mechanism in biology. Stud. Hist. Philos. Sci. Part C Stud. Hist. Philos. Biol. Biomed. Sci. 43, 152–163 (2012).

4. Asyali, M. H., Colak, D., Demirkaya, O. & Inan, M. S. Gene Expression Profile Classification: A Review. Curr. Bioinforma. 1, 55–73 (2006).

5. Reis-Filho, J. S. & Pusztai, L. Gene expression profiling in breast cancer: classification, prognostication, and prediction. The Lancet 378, 1812–1823 (2011).

6. Adzhubei, I., Jordan, D. M. & Sunyaev, S. R. Predicting Functional Effect of Human Missense Mutations Using PolyPhen-2. Curr. Protoc. Hum. Genet. 76, 7.20.1-7.20.41 (2013).

7. Meyer, M. J. et al. mutation3D: cancer gene prediction through atomic clustering of coding variants in the structural proteome. Hum. Mutat. 37, 447–456 (2016).

8. Shukla, K. et al. Machine Learning of Three-Dimensional Protein Structures to Predict the Functional Impacts of Genome Variation. J. Chem. Inf. Model. 64, 5328–5343 (2024).

9. Shukla, K., Wang, Y., Spanheimer, P. M. & Brunk, E. AI-Driven Variant Annotation for Precision Oncology in Breast Cancer. 2025.03.14.643357 Preprint at 10.1101/2025.03.14.643357 (2025).

10. Tsherniak, A. et al. Defining a Cancer Dependency Map. Cell 170, 564-576.e16 (2017).

11. Barretina, J. et al. The Cancer Cell Line Encyclopedia enables predictive modelling of anticancer drug sensitivity. Nature 483, 603–607 (2012).

12. Ghandi, M. et al. Next-generation characterization of the Cancer Cell Line Encyclopedia. Nature 569, 503–508 (2019).

13. Basu, A. et al. An interactive resource to identify cancer genetic and lineage dependencies targeted by small molecules. Cell 154, 1151–1161 (2013).

14. Abramson, J. et al. Accurate structure prediction of biomolecular interactions with AlphaFold 3. Nature 630, 493–500 (2024).

15. Berman, H. M. et al. The Protein Data Bank. Nucleic Acids Res. 28, 235–242 (2000).

16. Ursu, O. et al. Massively parallel phenotyping of coding variants in cancer with Perturb-seq. Nat. Biotechnol. 40, 896–905 (2022).

17. Huang, A. C. et al. X-Atlas/Orion: Genome-wide Perturb-seq Datasets via a Scalable Fix-Cryopreserve Platform for Training Dose-Dependent Biological Foundation Models. 2025.06.11.659105 Preprint at 10.1101/2025.06.11.659105 (2025).

18. Zhang, J. et al. Tahoe-100M: A Giga-Scale Single-Cell Perturbation Atlas for Context-Dependent Gene Function and Cellular Modeling. 2025.02.20.639398 Preprint at 10.1101/2025.02.20.639398 (2025).

19. Ng, P. K.-S. et al. Systematic Functional Annotation of Somatic Mutations in Cancer. Cancer Cell 33, 450-462.e10 (2018).

20. Glusman, G. et al. Mapping genetic variations to three-dimensional protein structures to enhance variant interpretation: a proposed framework. Genome Med. 9, 113 (2017).

21. Pieper, U. et al. modbase, a database of annotated comparative protein structure models and associated resources. Nucleic Acids Res. 37, D347–D354 (2009).

22. Getis, A. & Ord, J. K. The Analysis of Spatial Association by Use of Distance Statistics. Geogr. Anal. 24, 189–206 (1992).

23. Subramanian, A. et al. Gene set enrichment analysis: A knowledge-based approach for interpreting genome-wide expression profiles. Proc. Natl. Acad. Sci. 102, 15545–15550 (2005).

24. Shi, J. & Walker, M. G. Gene Set Enrichment Analysis (GSEA) for Interpreting Gene Expression Profiles. Curr. Bioinforma. 2, 133–137 (2007).

25. Liberzon, A. et al. The Molecular Signatures Database (MSigDB) hallmark gene set collection. Cell Syst. 1, 417–425 (2015).

26. Yevshin, I., Sharipov, R., Valeev, T., Kel, A. & Kolpakov, F. GTRD: a database of transcription factor binding sites identified by ChIP-seq experiments. Nucleic Acids Res. 45, D61–D67 (2017).

27. Baird, L. & Yamamoto, M. The Molecular Mechanisms Regulating the KEAP1-NRF2 Pathway. Mol. Cell. Biol. 40, e00099–20 (2020).

28. Leinonen, H. M., Kansanen, E., Pölönen, P., Heinäniemi, M. & Levonen, A.-L. Dysregulation of the Keap1–Nrf2 pathway in cancer. Biochem. Soc. Trans. 43, 645–649 (2015).

29. Lo, S.-C., Li, X., Henzl, M. T., Beamer, L. J. & Hannink, M. Structure of the Keap1:Nrf2 interface provides mechanistic insight into Nrf2 signaling. EMBO J. 25, 3605–3617 (2006).

30. Taguchi, K. & Yamamoto, M. The KEAP1–NRF2 System in Cancer. Front. Oncol. 7, (2017).

31. Suzuki, T. & Yamamoto, M. Molecular basis of the Keap1–Nrf2 system. Free Radic. Biol. Med. 88, 93–100 (2015).

32. Hayes, J. D. & McMahon, M. NRF2 and KEAP1 mutations: permanent activation of an adaptive response in cancer. Trends Biochem. Sci. 34, 176–188 (2009).

33. Hast, B. E. et al. Cancer-Derived Mutations in KEAP1 Impair NRF2 Degradation but not Ubiquitination. Cancer Res. 74, 808–817 (2014).

34. Fowler, D. M. & Fields, S. Deep mutational scanning: a new style of protein science. Nat. Methods 11, 801–807 (2014).

35. Esposito, D. et al. MaveDB: an open-source platform to distribute and interpret data from multiplexed assays of variant effect. Genome Biol. 20, 223 (2019).

36. Rubin, A. F. et al. MaveDB 2024: a curated community database with over seven million variant effects from multiplexed functional assays. Genome Biol. 26, 13 (2025).

37. Alvarez, M. J. et al. Network-based inference of protein activity helps functionalize the genetic landscape of cancer. Nat. Genet. 48, 838–847 (2016).

38. Padi, M. & Quackenbush, J. Detecting phenotype-driven transitions in regulatory network structure. Npj Syst. Biol. Appl. 4, 16 (2018).

39. Ouyang, B. et al. AI-powered omics-based drug pair discovery for pyroptosis therapy targeting triple-negative breast cancer. Nat. Commun. 15, 7560 (2024).

40. Glass, K., Huttenhower, C., Quackenbush, J. & Yuan, G.-C. Passing Messages between Biological Networks to Refine Predicted Interactions. PLOS ONE 8, e64832 (2013).

41. Franceschini, A. et al. STRING v9.1: protein-protein interaction networks, with increased coverage and integration. Nucleic Acids Res. 41, D808–815 (2013).

42. Jensen, L. J. et al. STRING 8--a global view on proteins and their functional interactions in 630 organisms. Nucleic Acids Res. 37, D412–416 (2009).

43. Snel, B., Lehmann, G., Bork, P. & Huynen, M. A. STRING: a web-server to retrieve and display the repeatedly occurring neighbourhood of a gene. Nucleic Acids Res. 28, 3442–3444 (2000).

44. Szklarczyk, D. et al. The STRING database in 2023: protein-protein association networks and functional enrichment analyses for any sequenced genome of interest. Nucleic Acids Res. 51, D638–D646 (2023).

45. Szklarczyk, D. et al. The STRING database in 2021: customizable protein-protein networks, and functional characterization of user-uploaded gene/measurement sets. Nucleic Acids Res. 49, D605–D612 (2021).

46. Szklarczyk, D. et al. STRING v11: protein-protein association networks with increased coverage, supporting functional discovery in genome-wide experimental datasets. Nucleic Acids Res. 47, D607–D613 (2019).

47. Szklarczyk, D. et al. The STRING database in 2017: quality-controlled protein-protein association networks, made broadly accessible. Nucleic Acids Res. 45, D362–D368 (2017).

48. Szklarczyk, D. et al. STRING v10: protein-protein interaction networks, integrated over the tree of life. Nucleic Acids Res. 43, D447–452 (2015).

49. Szklarczyk, D. et al. The STRING database in 2011: functional interaction networks of proteins, globally integrated and scored. Nucleic Acids Res. 39, D561–568 (2011).

50. von Mering, C. et al. STRING 7--recent developments in the integration and prediction of protein interactions. Nucleic Acids Res. 35, D358–362 (2007).

51. von Mering, C. et al. STRING: known and predicted protein-protein associations, integrated and transferred across organisms. Nucleic Acids Res. 33, D433–437 (2005).

52. von Mering, C. et al. STRING: a database of predicted functional associations between proteins. Nucleic Acids Res. 31, 258–261 (2003).

53. Oughtred, R. et al. The BioGRID database: A comprehensive biomedical resource of curated protein, genetic, and chemical interactions. Protein Sci. Publ. Protein Soc. 30, 187–200 (2021).

54. Popescu, M.-C., Balas, V. E., Perescu-Popescu, L. & Mastorakis, N. Multilayer perceptron and neural networks. WSEAS Trans Cir Sys 8, 579–588 (2009).

55. Guan, N.-N. et al. Anticancer Drug Response Prediction in Cell Lines Using Weighted Graph Regularized Matrix Factorization. Mol. Ther. - Nucleic Acids 17, 164–174 (2019).

56. Wang, L., Li, X., Zhang, L. & Gao, Q. Improved anticancer drug response prediction in cell lines using matrix factorization with similarity regularization. BMC Cancer 17, 513 (2017).

57. Suphavilai, C., Bertrand, D. & Nagarajan, N. Predicting Cancer Drug Response using a Recommender System. Bioinformatics 34, 3907–3914 (2018).

58. Huang, L., Guo, Z., Wang, F. & Fu, L. KRAS mutation: from undruggable to druggable in cancer. Signal Transduct. Target. Ther. 6, 386 (2021).

59. Jančík, S., Drábek, J., Radzioch, D. & Hajdúch, M. Clinical Relevance of KRAS in Human Cancers. BioMed Res. Int. 2010, 150960 (2010).

60. McCormick, F. KRAS as a Therapeutic Target. Clin. Cancer Res. 21, 1797–1801 (2015).

61. González-Alonso, P. et al. The Hippo Pathway Transducers YAP1/TEAD Induce Acquired Resistance to Trastuzumab in HER2-Positive Breast Cancer. Cancers 12, 1108 (2020).

62. Masliantsev, K., Karayan-Tapon, L. & Guichet, P.-O. Hippo Signaling Pathway in Gliomas. Cells 10, 184 (2021).

63. Edwards, A. C. et al. TEAD Inhibition Overcomes YAP1/TAZ-Driven Primary and Acquired Resistance to KRASG12C Inhibitors. Cancer Res. 83, 4112–4129 (2023).

64. Yang, X. et al. Abstract 1931: Targeting YAP1/TEAD signaling re-sensitizes MAPK/ERK pathway inhibitors in KRAS-driven cancer cells. Cancer Res. 84, 1931 (2024).

65. Drosten, M. & Barbacid, M. Targeting the MAPK Pathway in KRAS-Driven Tumors. Cancer Cell 37, 543–550 (2020).

66. Paul, S. et al. Cooperation between the Hippo and MAPK pathway activation drives acquired resistance to TEAD inhibition. Nat. Commun. 16, 1743 (2025).

67. Huh, H. D., Kim, D. H.Jeong, H.-S. & Park, H. W. Regulation of TEAD Transcription Factors in Cancer Biology. Cells 8, 600 (2019).

68. Wang, P. et al. Mu-KRAS attenuates Hippo signaling pathway through PKCι to sustain the growth of pancreatic cancer. J. Cell. Physiol. 235, 408–420 (2020).

69. Cunningham, R. & Hansen, C. G. The Hippo pathway in cancer: YAP/TAZ and TEAD as therapeutic targets in cancer. Clin. Sci. 136, 197–222 (2022).

70. Heng, B. C. et al. An overview of signaling pathways regulating YAP/TAZ activity. Cell. Mol. Life Sci. CMLS 78, 497–512 (2020).

71. Moroishi, T. et al. A YAP/TAZ-induced feedback mechanism regulates Hippo pathway homeostasis. Genes Dev. 29, 1271–1284 (2015).

72. Piccolo, S., Dupont, S. & Cordenonsi, M. The Biology of YAP/TAZ: Hippo Signaling and Beyond. Physiol. Rev. 94, 1287–1312 (2014).

73. Rausch, V. & Hansen, C. G. The Hippo Pathway, YAP/TAZ, and the Plasma Membrane. Trends Cell Biol. 30, 32–48 (2020).

74. Varelas, X. The Hippo pathway effectors TAZ and YAP in development, homeostasis and disease. Development 141, 1614–1626 (2014).

75. Pobbati, A. V., Kumar, R., Rubin, B. P. & Hong, W. Therapeutic targeting of TEAD transcription factors in cancer. Trends Biochem. Sci. 48, 450–462 (2023).

76. Hagenbeek, T. J. et al. An allosteric pan-TEAD inhibitor blocks oncogenic YAP/TAZ signaling and overcomes KRAS G12C inhibitor resistance. Nat. Cancer 4, 812–828 (2023).

77. Kumar, R. et al. Discovery of Pan-TEAD Inhibitors That Disrupt YAP-TEAD Interaction as a Potential Therapy for Gastric Cancers and Mutant KRAS and EGFR Lung Cancers. ACS Med. Chem. Lett. 17, 649–661 (2026).

78. Weinstein, J. N. et al. The Cancer Genome Atlas Pan-Cancer Analysis Project. Nat. Genet. 45, 1113–1120 (2013).

79. Landrum, M. J. et al. ClinVar: public archive of relationships among sequence variation and human phenotype. Nucleic Acids Res. 42, D980–985 (2014).

80. Suehnholz, S. P. et al. Quantifying the Expanding Landscape of Clinical Actionability for Patients with Cancer. Cancer Discov. 14, 49–65 (2024).

81. Chakravarty, D. et al. OncoKB: A Precision Oncology Knowledge Base. JCO Precis. Oncol. 1–16 (2017) doi:10.1200/PO.17.00011.

82. Mazzocchi, F. Complexity and the reductionism–holism debate in systems biology. WIREs Syst. Biol. Med. 4, 413–427 (2012).

83. Brigandt, I. & Love, A. Reductionism in Biology. in The Stanford Encyclopedia of Philosophy (eds. Zalta, E. N. & Nodelman, U.) (Metaphysics Research Lab, Stanford University, 2012).

84. Glick, D. Methods of Biochemical Analysis. (John Wiley & Sons, 2009).

85. Michal, G. & Schomburg, D. Biochemical Pathways: An Atlas of Biochemistry and Molecular Biology. (John Wiley & Sons, 2012).

86. Satyanarayana, U. Biochemistry. (Elsevier Health Sciences, 2013).

87. Schaffner, K. F. Reductionism in Biology: Prospects and Problems. PSA Proc. Bienn. Meet. Philos. Sci. Assoc. 1974, 612–632 (1974).

88. Sekhon, B. S. Chemical Biology: Past, Present and Future. Curr. Chem. Biol. 2, 278–311 (2008).

89. Gutteridge, A. & Thornton, J. M. Understanding nature’s catalytic toolkit. Trends Biochem. Sci. 30, 622–629 (2005).

90. Holliday, G. L., Almonacid, D. E., Mitchell, J. B. O. & Thornton, J. M. The Chemistry of Protein Catalysis. J. Mol. Biol. 372, 1261–1277 (2007).

91. Fersht, A. Structure and Mechanism in Protein Science: A Guide to Enzyme Catalysis and Protein Folding. (kMacmillan, 1999).

92. Adams, J. A. Kinetic and Catalytic Mechanisms of Protein Kinases. Chem. Rev. 101, 2271–2290 (2001).

93. Levy, Y., Wolynes, P. G. & Onuchic, J. N. Protein topology determines binding mechanism. Proc. Natl. Acad. Sci. 101, 511–516 (2004).

94. Stites, W. E. Protein™Protein Interactions: Interface Structure, Binding Thermodynamics, and Mutational Analysis. Chem. Rev. 97, 1233–1250 (1997).

95. Reichmann, D., Rahat, O., Cohen, M., Neuvirth, H. & Schreiber, G. The molecular architecture of protein–protein binding sites. Curr. Opin. Struct. Biol. 17, 67–76 (2007).

96. Cao, L. et al. Design of protein-binding proteins from the target structure alone. Nature 605, 551–560 (2022).

97. Petsko, G. A. & Ringe, D. Protein Structure and Function. (New Science Press, 2004).

98. Pawson, T. & Nash, P. Assembly of Cell Regulatory Systems Through Protein Interaction Domains. Science 300, 445–452 (2003).

99. Jr, D. E. K. Protein Shape and Biological Control. Scientific American https://www.scientificamerican.com/article/protein-shape-and-biological-contro/ (1973).

100. Cohen, P. The structure and regulation of protein phosphatases. Annu. Rev. Biochem. 58, 453–508 (1989).

101. Tomkins, G. M., Yielding, K. L., Talal, N. & Curran, J. F. Protein Structure and Biological Regulation. Cold Spring Harb. Symp. Quant. Biol. 28, 461–471 (1963).

102. Brändén, C.-I. & Tooze, J. Introduction to Protein Structure. (Taylor & Francis, 1999).

103. Rose, M. R. & Oakley, T. H. The new biology: beyond the Modern Synthesis. Biol. Direct 2, 30 (2007).

104. Amaral, L. A. N. & Ottino, J. M. Complex systems and networks: challenges and opportunities for chemical and biological engineers. Chem. Eng. Sci. 59, 1653–1666 (2004).

105. Kampis, G. Self-Modifying Systems in Biology and Cognitive Science: A New Framework for Dynamics, Information and Complexity. (Elsevier, 2013).

106. Loewe, L. A framework for evolutionary systems biology. BMC Syst. Biol. 3, 27 (2009).

107. Krakauer, D. C. et al. The challenges and scope of theoretical biology. J. Theor. Biol. 276, 269–276 (2011).

108. Chan, E. Y. Advances in sequencing technology. Mutat. Res. - Fundam. Mol. Mech. Mutagen. 573, 13–40 (2005).

109. Buermans, H. P. J. & den Dunnen, J. T. Next generation sequencing technology: Advances and applications. Biochim. Biophys. Acta BBA - Mol. Basis Dis. 1842, 1932–1941 (2014).

110. Pareek, C. S., Smoczynski, R. & Tretyn, A. Sequencing technologies and genome sequencing. J. Appl. Genet. 52, 413–435 (2011).

111. Satam, H. et al. Next-Generation Sequencing Technology: Current Trends and Advancements. Biology 12, 997 (2023).

112. Kherlopian, A. R. et al. A review of imaging techniques for systems biology. BMC Syst. Biol. 2, 74 (2008).

113. Weissleder, R. & Nahrendorf, M. Advancing biomedical imaging. Proc. Natl. Acad. Sci. 112, 14424–14428 (2015).

114. Dai, X. & Shen, L. Advances and Trends in Omics Technology Development. Front. Med. 9, (2022).

115. Baysoy, A., Bai, Z., Satija, R. & Fan, R. The technological landscape and applications of single-cell multi-omics. Nat. Rev. Mol. Cell Biol. 24, 695–713 (2023).

116. Radivojac, P. et al. A large-scale evaluation of computational protein function prediction. Nat. Methods 10, 221–227 (2013).

117. Santolini, M. & Barabási, A.-L. Predicting perturbation patterns from the topology of biological networks. Proc. Natl. Acad. Sci. 115, E6375–E6383 (2018).

118. Molinelli, E. J. et al. Perturbation Biology: Inferring Signaling Networks in Cellular Systems. PLOS Comput. Biol. 9, e1003290 (2013).

119. Tam, V. et al. Benefits and limitations of genome-wide association studies. Nat. Rev. Genet. 20, 467–484 (2019).

120. Baranzini, S. E. et al. Genome-wide association analysis of susceptibility and clinical phenotype in multiple sclerosis. Hum. Mol. Genet. 18, 767–778 (2009).

121. Sud, A., Kinnersley, B. & Houlston, R. S. Genome-wide association studies of cancer: current insights and future perspectives. Nat. Rev. Cancer 17, 692–704 (2017).

122. Kumar, P., Henikoff, S. & Ng, P. C. Predicting the effects of coding non-synonymous variants on protein function using the SIFT algorithm. Nat. Protoc. 4, 1073–1081 (2009).

123. Araya, C. L. & Fowler, D. M. Deep mutational scanning: assessing protein function on a massive scale. Trends Biotechnol. 29, 435–442 (2011).

124. Ng, P. C. & Henikoff, S. Predicting the Effects of Amino Acid Substitutions on Protein Function. Annu. Rev. Genomics Hum. Genet. 7, 61–80 (2006).

125. Carninci, P. et al. The Transcriptional Landscape of the Mammalian Genome. Science 309, 1559–1563 (2005).

126. Braxton, A. M. et al. 3D genomic mapping reveals multifocality of human pancreatic precancers. Nature 629, 679–687 (2024).

127. Ashburner, M. et al. Gene Ontology: tool for the unification of biology. Nat. Genet. 25, 25–29 (2000).

128. The Gene Ontology Consortium. The Gene Ontology knowledgebase in 2026. Nucleic Acids Res. 54, D1779–D1792 (2026).

129. Kotlyar, M., Pastrello, C., Sheahan, N. & Jurisica, I. Integrated interactions database: tissue-specific view of the human and model organism interactomes. Nucleic Acids Res. 44, D536–541 (2016).

130. Swenson, N., Krishnapriyan, A. S., Buluc, A., Morozov, D. & Yelick, K. PersGNN: Applying Topological Data Analysis and Geometric Deep Learning to Structure-Based Protein Function Prediction. Preprint at 10.48550/arXiv.2010.16027 (2020).

131. Ma, J. et al. Using deep learning to model the hierarchical structure and function of a cell. Nat. Methods 15, 290–298 (2018).

132. Adamovich, A. I. et al. The functional impact of BRCA1 BRCT domain variants using multiplexed DNA double-strand break repair assays. Am. J. Hum. Genet. 109, 618–630 (2022).

133. Kato, S. et al. Understanding the function–structure and function–mutation relationships of p53 tumor suppressor protein by high-resolution missense mutation analysis. Proc. Natl. Acad. Sci. 100, 8424–8429 (2003).

134. Weng, C., Faure, A. J., Escobedo, A. & Lehner, B. The energetic and allosteric landscape for KRAS inhibition. Nature 626, 643–652 (2024).

135. Wang, Y.-X. & Zhang, Y.-J. Nonnegative Matrix Factorization: A Comprehensive Review. IEEE Trans. Knowl. Data Eng. 25, 1336–1353 (2013).

136. Multidimensional Scaling. in Measurement, Judgment and Decision Making 179–250 (Academic Press, 1998). doi:10.1016/B978-012099975-0.50005-1.

137. Sinaga, K. P. & Yang, M.-S. Unsupervised K-Means Clustering Algorithm. IEEE Access 8, 80716–80727 (2020).

138. Ahmad, S. et al. The UniProt website API: facilitating programmatic access to protein knowledge. Nucleic Acids Res. 53, W547–W553 (2025).

139. Getis, A. & Aldstadt, J. Constructing the Spatial Weights Matrix Using a Local Statistic. Geogr. Anal. 36, 90–104 (2004).

